# Model-Based Feature Selection and Clustering of Rna-Seq Data for Unsupervised Subtype Discovery

**DOI:** 10.1101/2020.05.23.111799

**Authors:** David K. Lim, Naim U. Rashid, Joseph G. Ibrahim

## Abstract

Clustering is a form of unsupervised learning that aims to uncover latent groups within data based on similarity across a set of features. A common application of this in biomedical research is in delineating novel cancer subtypes from patient gene expression data, given a set of informative genes. However, it is typically unknown *a priori* what genes may be informative in discriminating between clusters, and what the optimal number of clusters are. Few methods exist for performing unsupervised clustering of RNA-seq samples, and none currently adjust for between-sample global normalization factors, select cluster-discriminatory genes, or account for potential confounding variables during clustering. To address these issues, we propose the Feature Selection and Clustering of RNA-seq (FSCseq): a model-based clustering algorithm that utilizes a finite mixture of regression (FMR) model and utilized the quadratic penalty method with a SCAD penalty. The maximization is done by a penalized Classification EM algorithm, allowing us to include normalization factors and confounders in our modeling framework. Given the fitted model, our framework allows for subtype prediction in new patients via posterior probabilities of cluster membership. Based on simulations and real data analysis, we show the advantages of our method relative to competing approaches.

## 1. Introduction

Clustering is a form of unsupervised learning that aims to uncover latent groups across subjects based on their similarity with respect to a common set of features. A common application of clustering is in the discovery of novel molecular subtypes in cancer based upon the patients’ subject-specific RNA sequencing (RNA-seq) gene expression profiles. Many studies have demonstrated the clinical utility of such molecular subtypes, where significant differences in patient prognosis and treatment response between subtypes have been observed (Perou et al., 2000; Chia et al., 2012; Mao et al., 2017).

However, few methods are able to cluster samples whose genes (features) are measured via RNA-seq, a common platform for measuring gene expression based upon high-throughput sequencing (Mo et al., 2013; Li et al., 2018). Gene expression in RNA-seq studies is typically quantified in terms of gene-level read counts, defined by the number of sequencing reads mapping to protein coding regions in a particular gene following sequence alignment. Gene expression quantitation software such as `RSEM` and `Salmon` can further account for for multimapped reads, or sequences that map to multiple regions of the genome (Li and Dewey, 2011; Patro et al., 2017), in addition to other technical biases in quantifying gene-level read counts. Such counts are often overdispersed, and are therefore commonly modeled by the negative binomial distribution (Robinson, McCarthy and Smyth, 2009; Love, Huber and Anders, 2014). This is in contrast to Gaussian distributional assumptions made by earlier methods designed for clustering subjects with microarray-based gene expression measurements, such as mclust (Scrucca et al., 2016).

This shift in distributional assumption has practical implications. For example, between-sample variation in technical factors such as sequencing depth can cause global shifts in sample-specific read count distributions, necessitating the use of between-sample correction methods (Li et al., 2015). However, due to the count nature of RNA-seq data, these correction factors are often directly incorporated into statistical modeling procedures, for example, as offset terms in negative binomial regression-based differential expression methods (Robinson, McCarthy and Smyth, 2009; Love, Huber and Anders, 2014). Correction for other confounders, such as batch effects (Leek et al., 2010) or specific clinical variables, is done by including the relevant factors as regression covariates. However, to the best of our knowledge, no analogous approach exists to correct for such factors in RNA-seq read count-based clustering analyses. Therefore, an approach that is able to correct for such factors may have significant utility in improving clustering performance.

Furthermore, a large number of genes is often available for clustering subjects (typically 20,000 or more), and it is often unknown *a priori* which subset of genes may best discriminate between latent clusters. Utilizing all available genes for clustering may negatively impact the accuracy and usefulness of derived clusters (Tritchler, Parkhomenko and Beyene, 2009), cause methods to underestimate the number of latent groups (Pan and Shen, 2007), and add significantly to computation time. As a result, it is common practice to filter a large proportion of genes prior to clustering, for example, by removing low variance genes with respect to their median absolute deviation (MAD) across samples (Chung et al., 2008; Wu et al., 2013). However, it is often unclear how many genes should be kept to ensure optimal clustering performance, as a poor threshold can decrease power of detecting differentially expressed genes (Bourgon, Gentleman and Huber, 2010). Lastly, the identification of such cluster-discriminatory genes may improve biological understanding of identified clusters via gene set enrichment analyses (Subramanian et al., 2005), and may aid in the development of future subtype classification methods (Golub et al., 1999).

Current methods that seek to cluster RNA-seq data have several limitations with respect to the issues mentioned above. We divide these methods into three general classes: transformation-based, count-based, and non-parametric clustering methods. Transformation-based methods first apply a variance-stabilizing transformation to RNA-seq gene read counts, allowing for the application of clustering methods developed for microarray data (Zwiener, Frisch and Binder, 2014) by assuming the transformed data are distributed approximately Gaussian. However, prior work has shown that the choice of transformation can greatly influence clustering performance, as the quality of this Gaussian approximation varies by condition, and the optimal choice may not be readily apparent (Noel-MacDonnell et al., 2018). Examples of such transformations include the log, variance stabilizing (Anders and Huber, 2010), and regularized log (rlog) (Love, Huber and Anders, 2014) transforms. The second class of methods directly models gene-level read counts assuming some type of count distribution. One such method is `iCluster+` (Mo et al., 2013), which assumes a Poisson distribution on the gene read counts. `iCluster+` also performs feature selection by inducing sparsity via the lasso penalty, but ignores potential overdispersion, or extra-Poisson variation, in counts. NBMB (Li et al., 2018) accounts for overdispersion via the negative-binomial distribution, but cannot identify cluster-discriminatory genes. In addition, neither method is able to adjust for confounding factors, such as batch effects or differences in sequencing depth. The third class of methods includes non-parametric clustering approaches such as the average-linkage hierarchical clustering (HC) and K-medoids (KM) (Jaskowiak, Costa and Campello, 2018). However, these algorithms cannot perform feature selection, nor can they adjust for batch effects or differences in sequencing depth. Lastly, none of these methods can be directly utilized to predict discovered subtypes in new subjects, limiting their direct translational utility.

To address these issues, we propose FSCseq (Feature Selection and Clustering of RNA-seq): a penalized finite mixture regression model that assumes negative-binomially distributed mixture components. In addition to directly clustering samples with respect to RNA-seq gene read counts, FSCseq also performs simultaneous feature selection to select cluster-discriminatory genes during the clustering process. We achieve this by imposing a fusion penalty on the differences in estimated means within each gene, simultaneously shrinking the estimates closer together while inducing sparsity on the differences across clusters. By modeling cluster means with weighted negative binomial regression models, FSCseq can directly adjust for differences in sequencing depth through the inclusion of size factors in a manner similar to common differential gene expression methods. Our method can also correct for potential confounders, such as batch effects, by including them as regression covariates. We demonstrate the utility of these features in improving clustering performance via simulation studies and a real data application to a TCGA breast cancer RNA-seq dataset. Lastly, we illustrate how FSC-seq performs subtype prediction in new samples, given a new set of gene expression profiles and a prior fitted FSCseq object. In all, we address critically lacking features in clustering samples whose gene expression profiles are measured via RNA-seq, and demonstrate improved performance with respect to existing methods over a range of conditions.

## 2. Methods

The main utility of FSCseq is two-fold, where we perform 1) unsupervised clustering of subjects based on subject-specific RNA-seq gene expression profiles, and 2) simultaneous selection of cluster-discriminatory genes. Our modeling approach can correct for sources of technical variation like sequencing depth, adjust for effects of potential confounders during clustering such as batch effects, and directly predict discovered subtypes in new patients based upon a previously fitted FSCseq model. We describe our model implementation below, and a visualization of the FSCseq workflow can be found in Section A of the Supplementary Material [Lim, 2020].

### 2.1. Model likelihood

Gene-level RNA-seq read counts are utilized as the basis of our model, which can be obtained from RNA-seq gene expression quantification software such as `Salmon` (Patro et al., 2017). Such approaches account for read multi-mapping and other biases and, as a result, often provide non-integer “expected read counts” adjusting for these factors. We round these expected count values to integers, similar to previous approaches (Love, Huber and Anders, 2014). Let **y** denote an *n × G* matrix of *n* subjects (or “samples”) and *G* genes (or “features”), with each element *y_ij_* denoting the read count of the *j^th^* gene for the *i^th^* subject, where *i* = 1*, … , n* and *j* = 1*, … , G*.

We assume that the RNA-seq gene expression profile of subject *i*, ***y***_*i*_ = (*y_i_*_1_*, … , y_iG_*), can be modeled by a mixture of *K G*-dimensional multivariate negative binomial regression models. The likelihood of this mixture model may be written as

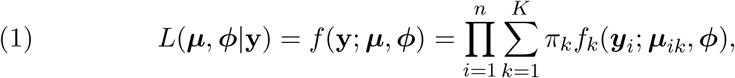

where *K* is the “order” of the mixture model (i.e. the number of assumed clusters in the data), *π_k_* is the mixture proportion pertaining to cluster *k*, and 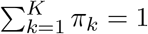. We denote *μ_ijk_* as the mean of the *j^th^* gene for the *i^th^* subject in the *k^th^* cluster, after adjusting for subject-specific sequencing depth, and we denote *ϕ_j_* as the gene-level dispersion parameter pertaining to gene *j*. Then, ***μ***_*ik*_ = (*μ_i_*_1*k*_, … , μ_*iGk*_) is the mean expression profile pertaining to sample *i* and cluster *k*, and ***ϕ*** = (*ϕ*_1_*, … , ϕ_G_*) is a vector of the *G* gene-level dispersion parameters.

We utilize a gene-gene independence assumption, which has been successfully implemented in prior RNA-seq read count-based clustering methods (Si et al., 2013; Mo et al., 2013; Li et al., 2018), as a reasonable approximation to simplify (1). Under this assumption, 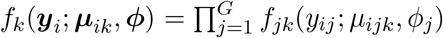, where *f_jk_*(*·*) is the negative binomial probability mass function, given by 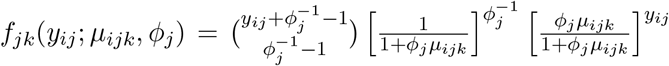, equivalent to the “NB2” parameterization by Hilbe (2009). Although the multivariate lognormal Poisson or multivariate negative binomial model may alternatively account for potential correlation between genes, the specification and estimation of such a covariance structure across genes is intractable in very high dimensions (Inouye et al., 2017).

For the *j^th^* gene, we may specify the link and variance functions with respect to *μ_ijk_* as

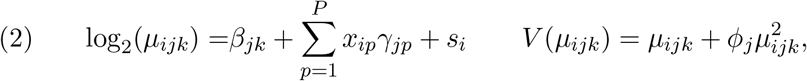

where *β_jk_* is the log base 2 (log_2_) mean read count for gene *j* and cluster *k*, *γ_jp_* is the effect of covariate *p* on the expression of gene *j*, and *s_i_* is the log_2_ scaled size factor of subject *i* calculated by `DESeq2` to correct for subject-specific differences in sequencing depth. In other words, we model the log_2_ mean of the *j^th^* gene for subjects in the *k^th^* cluster via a cluster-specific mean parameter (on the log_2_ scale) and a set of *P* subject-level covariates (*x_i_*_1_*, … , x_iP_*), while also accounting for sequencing depth. We allow covariate effects to be distinct across genes, similar to prior batch correction methods (Johnson, Li and Rabinovic, 2006; Leek, 2014), as, biologically, certain sets of genes may be affected differently by a particular covariate. Examples of such covariates may include technical factors such as batch, or potential demographic variables such as age or tumor grade. Through the inclusion of covariates in the model, we will later demonstrate FSCseq’s ability to adjust cluster mean estimates, thus improving clustering performance in the presence of such factors.

We note that in the given parametrization, we specify gene-level dispersion parameters *ϕ_j_* that are shared across the *K* clusters within gene *j* (Robinson, McCarthy and Smyth, 2009; Love, Huber and Anders, 2014). In contrast, NBMB (Li et al., 2018) specifies cluster-specific dispersion parameters ***ϕ***_*j*_ = (*ϕ_j_*_1_ *… , ϕ_jK_*) within each gene *j*. Conceptually, when *n* is small, the estimate of each *ϕ_jk_* may not be accurate due to the smaller average per-cluster sample size (Piegorsch, 1990; Al-Khasawn, 2010). In FSCseq, we utilize gene-level dispersions as the default in order to avoid this issue, and later show that the cluster-specific dispersion scheme by NBMB performs poorly in clustering analyses.

#### 2.1.1. Selection of cluster-discriminatory genes via fusion penalty

FSC-seq identifies cluster-discriminatory genes simultaneously during clustering. To do this, within a given gene *j*, we impose the SCAD penalty (Fan and Li, 2001) on the pairwise differences between log_2_ cluster means, in order to induce shrinkage on these differences and enforce equality of similar group means within gene *j*. That is, we introduce the penalty term

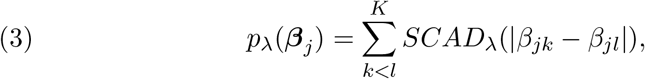

where ***β**_*j*_* = (*β_j_*_1_*, … , β_jK_*), *λ* is a tuning parameter to control the amount of penalty that is introduced, and *k* and *l* are cluster indices such that 1 *≤ k < l ≤ K*. The explicit form of the original SCAD penalty can be found in Section A1 of the Supplementary Material [Lim, 2020].

Then, we aim to maximize the penalized log-likelihood, which may be written as

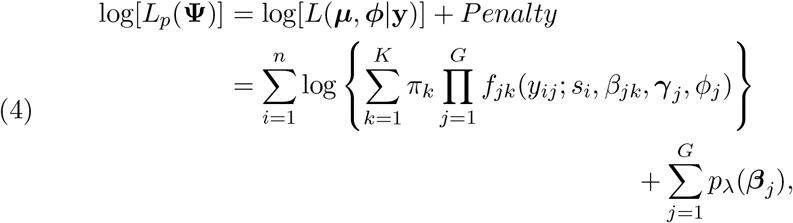

after incorporating the link function in (2) and the SCAD penalty. Here, **Ψ**= (***π**, **β**, **γ**, **ϕ***) pertains to the vector of model parameters, with ***π*** = (*π*_1_*, … , π_K_*), ***β*** = (***β***_1_*, … , **β**_G_*), and ***γ*** = (***γ***_1_*, … , **γ**_G_*). Here, ***γ**_*j*_* = (*γ_j_*_1_*, … , γ_jP_*) denotes the regression coefficients pertaining to the *P* covariates in gene *j*, *j* = 1*, … , G*. The first term of (4) is the log of the model likelihood in (1), with *μ_ijk_* replaced by (*s*_i_, β_jk_, **γ**_*j*_), since *μ_ijk_* is a function of (*s_i_, β_jk_, **γ**_*j*_*) by (2).

However, because the penalty is imposed on the differences in cluster log_2_ mean parameters in gene *j*, it is not separable in ***β**_*j*_*. As a result, the penalized log-likelihood may not converge to a stationary point during maximization (Friedman et al., 2007; Wu and Lange, 2008). Similar to Pan, Shen and Liu (2013), we address this by introducing a new variable *θ_j,kl_* = *β_jk_ − β_jl_* for each gene *j* = 1*, … , G* and each pair of clusters *k* and *l* such that 1 *≤ k < l ≤ K*. We then apply the quadratic penalty method (Nocedal and Wright, 2000) leveraging this new constraint. In particular, for a given gene *j*, let ***θ**_j_* be a *K × K* upper triangular matrix denoting the pairwise differences in ***β**_*j*_*, with entries (***θ**_j_*)_*kl*_ = *θ_j,kl_*. Then, under the quadratic penalty method, (3) can be rewritten as

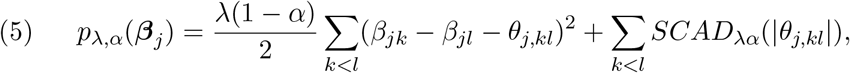

where the first term is the added quadratic penalty term as described by Nocedal and Wright (2000), pertaining to the constraint *θ_j,kl_* = *β_jk_ − β_jl_*. This first term in (5) is convex and differentiable, while the second term is non-smooth, but separable and convex, and so a coordinate-wise descent algorithm is guaranteed to converge to the global optimum (Tseng, 2001). In FSCseq, we utilize this property to implement a coordinate-wise descent algorithm (CDA) to estimate cluster log_2_ means and covariate effects within each given gene *j* = 1*, … , G* (See Section 2.2.2 for more details). We also introduce the hyperparameter *α* to control the balance between these two terms. Although similar in form, we note that this is not the traditional elastic net penalty (Zou and Hastie, 2005). Rather, this penalty is similar to the L1 penalty with the quadratic penalty method proposed by (Pan, Shen and Liu, 2013), with the L1 replaced by the SCAD penalty.

We also note that *β_jk_ −β_jl_ →* 0 as *θ_j,kl_ →* 0. However, similar to Pan, Shen and Liu (2013), we observe that *β_jk_ − β_jl_ − θ_j,kl_ →* 0 as *λ*(1 *− α*) *→ ∞*, but *β_jk_ − β_jl_* = 0 cannot be guaranteed. In particular, let 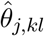 and 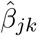 denote the estimates of *θ_j,kl_* and *β_jk_*, respectively. Then, although 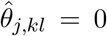 can be attained by the SCAD penalty, 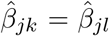 cannot be guaranteed for two distinct clusters *k* and *l*. This is because the term introduced by the quadratic penalty method (Nocedal and Wright, 2000) to enforce the constraint *θ_j,kl_* = *β_jk_ − β_jl_* is a ridge-like term, and thus does not perform strict thresholding, much like the classic L2 penalty (Hoerl and Kennard, 1970). Increasing the first term by increasing *λ*(1 *− α*) more closely approximates equality in the constraint, but this introduces bias in the estimation of ***β**_*j*_* (Pan, Shen and Liu, 2013).

In FSCseq, we address this by fusing clusters *k* and *l* together into one joint cluster for gene *j* if 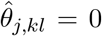, denoting the index of this new cluster as *k_*_*. Specifically, we estimate 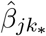 using data from both clusters *k* and *l* in gene *j*, and we strictly enforce the quadratic penalty method constraint by setting 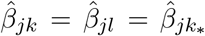. If all clusters are fused together in this way for a particular gene *j*, then gene *j* is determined to be “nondiscriminatory” across clusters. Otherwise, gene *j* is considered “cluster-discriminatory.” We call this penalty the “SCAD fusion penalty”. See Section 2.2 and Section A1 in the Supplementary Material for more details on this procedure [Lim, 2020].

We select optimal values of the three tuning parameters (*K*, *α*, and *λ*) by searching over a grid of possible values for *α* and *λ* with *K* fixed, and then repeat this procedure for every candidate value of *K*. We utilize the Bayesian Information Criterion (BIC) for selection (Schwarz, 1978). Details of this tuning parameter selection procedure is given in Section 2.2.6.

### 2.2. Computation

We maximize the penalized log-likelihood from (4) via a Classification Expectation-Maximization (CEM) algorithm with simulated annealing (Celeux and Govaert, 1992) to reduce the likelihood of converging to a local maximum. To reduce computational overhead per iteration, we also apply “mini-batching” of genes. That is, we update the estimates of parameters pertaining to a random subset (or a ‘mini-batch’) of the genes in the M-step, and then perform the E-step after each ‘minibatched’ M-step. This is in contrast to the standard EM algorithm, where we would only perform the E-step after estimating parameters for all genes in the M-step. Thus, mini-batching allows the E-step to be updated more quickly with M-step updates from a subset of the parameter space, similar to the multicycle ECM algorithm (Meng and Rubin, 1993) which has been shown to decrease overall computation time (Meng, 1994). We show later that this mini-batching approach works well in our simulation and real data analyses.

The complete data penalized log-likelihood, which corresponds to the penalized log-likelihood in (4), can be written as

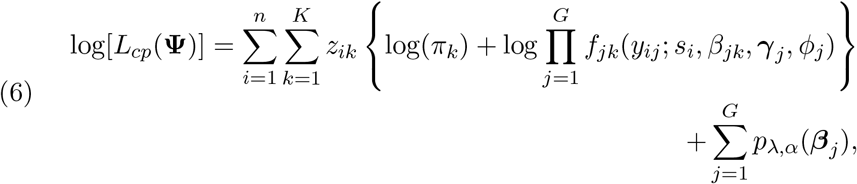

where *z_ik_* = *I*(*z_i_* = *k*) is the indicator of whether subject *i* belongs to cluster *k*. In this context, *z_ik_* is a latent variable denoting cluster membership of sample *i* in cluster *k*. In the standard EM algorithm, the so-called “Q-function” is then calculated as the conditional expectation of log[*L_cp_*(**Ψ**)], given the current parameter estimates and observed data (McLachlan and Krishnan, 2008; Garcia, Ibrahim and Zhu, 2009). This expectation is calculated in the E-step, and parameter estimates are updated in the M-step by maximizing the Q-function.

The rest of this section is organized as follows. To simplify the presentation of our method, we first describe the general E and M-steps for our algorithm without mini-batching, given some fixed values of *K*, *α*, and *λ*, in Sections 2.2.1 and 2.2.2. Then, we describe the modifications of these steps to support gene mini-batching in Section 2.2.3. We then show how the FSCseq algorithm is initialized in Section 2.2.4, outline convergence criteria in Section 2.2.5, and describe our procedure for selecting the optimal values of *K*, *α* and *λ* in Section 2.2.6. We conclude this section by describing a method for subtype prediction using the converged FSCseq model in Section 2.2.7.

#### 2.2.1. E-Step

We first describe the E-step with no mini-batching of genes. Let *m* denote the current EM iteration, 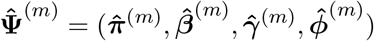 denote the estimates of (***π**, **β**, **γ**, **ϕ***) in the current iteration, and 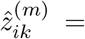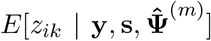 denote the conditional expectation of *z_ik_*, given the observed data and the current parameter estimates, where **s**= (*s*_1_*, … , s_n_*) is the vector of fixed size factor offsets calculated *a priori* by `DESeq2` (Love, Huber and Anders, 2014). Given the form of (6), evaluation of the Q-function requires only the calculation of 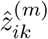, which can be interpreted as the posterior probability of subject *i* belonging to cluster *k* given the parameter estimates at the *m^th^* EM iteration. Thus, in the standard EM algorithm, the *m^th^* E-step reduces to computing 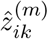 for *i* = 1*, … , n* and *k* = 1*, … , K* by the following expression:

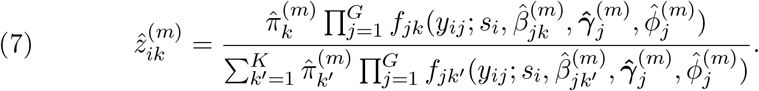

These posterior probabilities may then be used as cluster-specific observation weights in the M-step. Following convergence, it is also common to assign cluster membership to a subject by determining the cluster with the maximum posterior probability in that subject. That is, we can assign subject *i* to the cluster corresponding to arg max_*k*_ 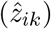 at convergence. We note from (7) that the contribution of cluster-nondiscriminatory genes cancel out in the posterior probability calculations.

However, the standard EM algorithm is known for its susceptibility to converge to a local optimum, rather than the global optimum (Dellaert, 2002). To reduce the likelihood of convergence to a local optimum, we modify the E-step in a manner similar to the Classification EM (CEM) algorithm with simulated annealing (Celeux and Govaert, 1990; Si et al., 2013). In the original CEM with simulated annealing algorithm (Celeux and Govaert, 1990), the E-step update in (7) is modified (denoted as the “AE-step”), and an additional sampling step (denoted as the “C-step”) is added after the AE-step.

In the AE-step, a “temperature” term is incorporated into the E-step update, and is gradually decreased (or “annealed”) such that the values of 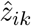 approach 0 or 1. The AE-step update corresponding to (7) is written as

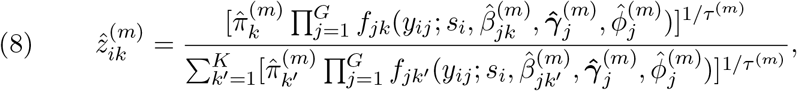

where *τ* ^(*m*)^ is the temperature of annealing in the *m^th^* AE-step iteration. As 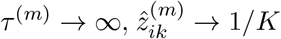, maximizing the entropy of the partition. As *τ* ^(*m*)^ *→* 0, we approach a hard partition for cluster membership (Rose, 1998) because the elements of the vector of posterior probabilities 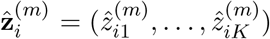 tend to 0 or 1, and 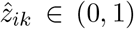 with 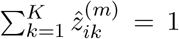 for all subjects *i* = 1*, … , n*. Traditionally, one allows *τ* ^(*m*)^ 0 as *m* at a pre-specified annealing rate *r* such that 0 *< r <* 1 and *τ* ^(*m*+1)^ = *rτ* ^(*m*)^. In this way, the AE-step modification allows the CEM to avoid a local optimum by gradually reducing the stochasticity introduced by the sampling in the C-step.

In the C-step, a sample **c**_*i*_ = (*c_i_*_1_*, … , c_iK_*) is drawn from the multinomial distribution with probabilities equal to 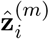 from (8). Then, we set 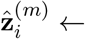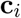, such that the 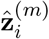 now specify a hard classification of sample *i*, i.e. for a given sample 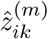 is now exactly 1 for one value of *k*, and 0 for all other values of *k*. This is repeated for all samples *i* = 1*, … , n*, yielding a hard classification partition of samples.

The combination of sampling in the C-step and the inclusion of annealing in the AE-step together helps the algorithm escape potential local optima. This is achieved by initializing *τ* ^(*m*)^ to be large in the first iterations, such that the 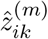 from the AE-step are close to 1*/K* for all *i* and *k*. Then, the probability of yielding a random partition of subjects in the C-step is high in the first iterations. By decreasing the value of *τ* ^(*m*)^, the probability of random partitioning approaches 0. Thus, a transition corresponding to a decrease in the objective function can be accepted with non-zero probability, and this probability approaches 0 as the algorithm proceeds (Celeux and Govaert, 1992). This results in the useful property for the CEM of not terminating when reaching the first local optimum.

Following the guidelines by Celeux and Govaert (1992) and Rose (1998), we set the annealing rate *r* = 0.9, similar to other methods (Si et al., 2013; Li et al., 2018). It is suggested that the entropy should be sufficiently large in early iterations, eventually shrinking to 0 in later iterations (Klein and Dubes, 1989; van Laarhoven and Aarts, 1987; Rose, 1998). Additionally, if the initial temperature *τ* ^(1)^ is too small, annealing may terminate at a suboptimal solution (Celeux and Govaert, 1992). We address this by setting the value of *τ* ^(1)^ = *G*. We find that this works well in our simulations and real data analyses.

#### 2.2.2. M-step

In the M-step, we maximize the Q-function, given the current E-step values 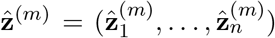 and some specified values of *α* and *λ*, updating the current estimates of the model parameters. From (6), the estimation of *π_k_* is separable from the remaining model parameters, and the corresponding updates for 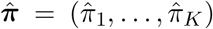 are given by 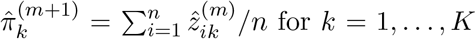. We also observe that the updates of the remaining model parameters are separable across genes. That is, we may obtain corresponding estimates 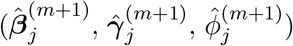 for each gene *j* separately of all other genes at each M-step, as these parameters are not shared across genes. This observation also facilitates the mini-batching approach described in Section 2.2.3. In this section, we first outline the M-step without mini-batching.

Specifically, for a given gene *j*, we utilize Iteratively Reweighted Least Squares (IRLS) with an embedded Coordinate Descent Algorithm (CDA) that updates 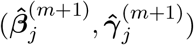, given specified values of 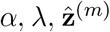, and 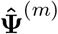. The update scheme for a particular gene *j* consists of two nested loops: an inner loop corresponding to the CDA and an outer loop corresponding to the IRLS. The IRLS weights and working responses are updated in the outer loop, and fed into the inner loop to update the parameter estimates in a coordinate-descent fashion. After convergence of IRLS, we estimate 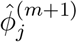 using Newton-Raphson. The entire M-step procedure is outlined in Algorithm 1.

##### Algorithm 1 M-step

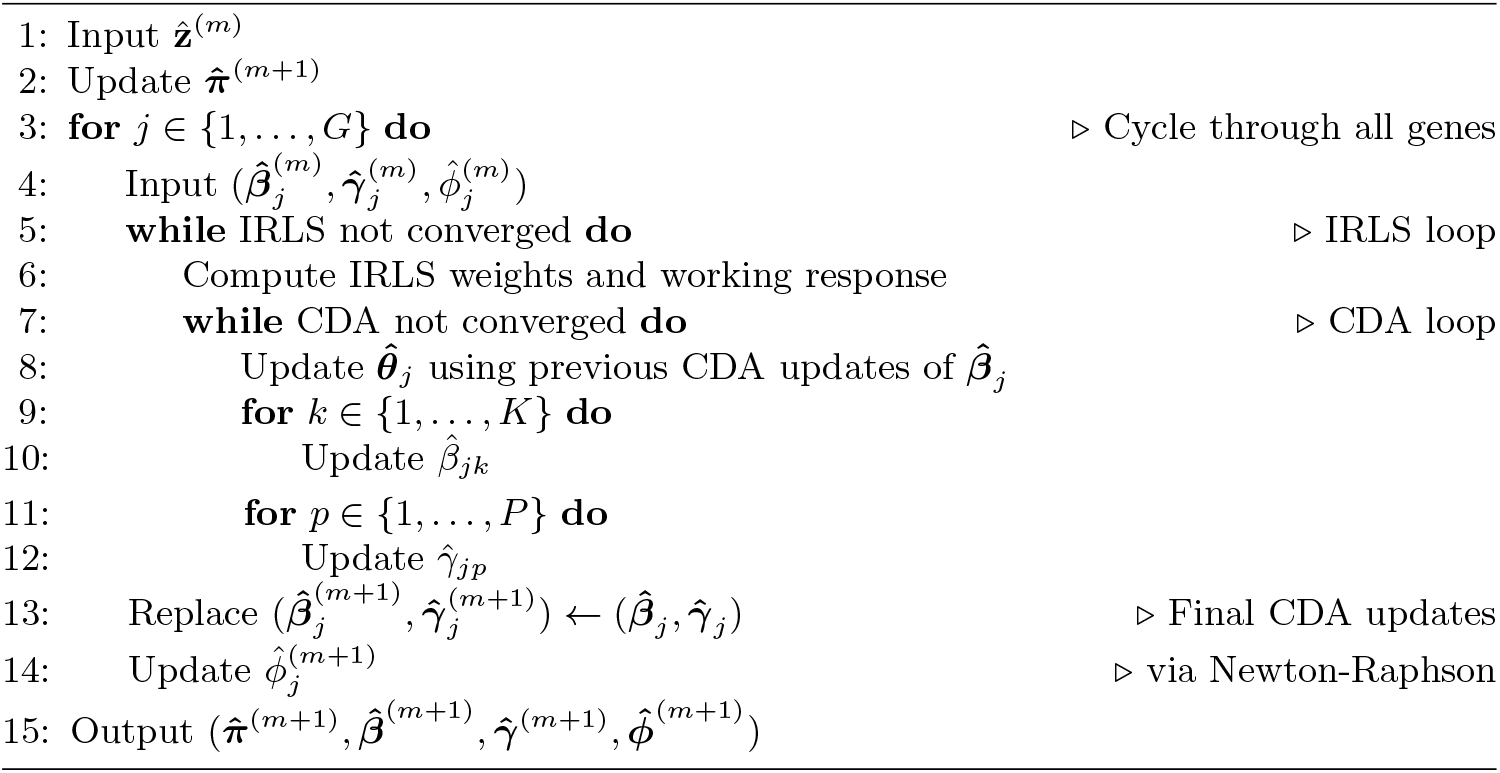

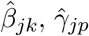, and 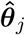 denote the CDA updates of *β_jk_*, *γ_jp_*, and ***θ**_j_*, respectively, and 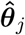 facilitates coordinate-wise descent updates of 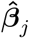. Notably, the fusion SCAD penalty is imposed on the updates of 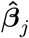, but updates of all other parameters are not penalized. Additional details of our CDA within IRLS implementation, including the forms of IRLS weights and working response updates, as well as the closed-form CDA updates pertaining to 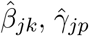, and 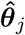 are given in Sections A2 and A3 of the Supplementary Material [Lim, 2020].

#### 2.2.3. Mini-batching of genes

In order to make our algorithm more scalable to large datasets and to speed up the computation time, we mini-batch genes in the M-step, such that the parameters pertaining to only a subset of genes are estimated during a particular M-step iteration (Neal and Hinton, 1998). Mini-batching of genes is done by randomly drawing a prespecified proportion of the genes whose parameters are updated in that M-step iteration. By default, we set the mini-batch size at each iteration to *G/*5, or 20% of the total genes. Specifically, in the M-step, we run Algorithm 1 on the pre-specified subset of genes corresponding to the current mini-batch. That is, we update 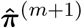 as before, but we update 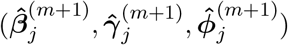 only if gene *j* is in the mini-batch for that M step iteration. This is straightforward to implement due to the separability across genes in parameter updates. If gene *j* is not in the mini-batch for the current M-step, current estimates are replaced with the previous iteration’s estimates, i.e. 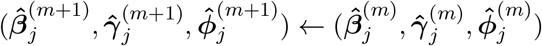.

The E-step calculations are then done as in Section 2.2.1 using the parameter estimates of all genes, with the updated values pertaining to the included genes of the mini-batch. Although mini-batching requires more CEM iterations to converge, we found that it yields slightly faster convergence than updating all genes at every M-step.

#### 2.2.4. Initialization

For each candidate value of *K*, we initialize our algorithm with very small penalty (small values of *α* and *λ*) from two sets of informative initial cluster labels (from naïve K-means and hierarchical clustering), and one set of random initial cluster labels drawn from the multinomial distribution with equal probabilities for the *K* clusters. We run the CEM with mini-batching on these 3 initializations until convergence, and select the converged model that yields the lowest BIC. Then, we run the standard EM algorithm from this selected starting point to convergence. This is similar to the “xCEM-EM” strategy proposed by Biernacki, Celeux and Govaert (2003), using the BIC for selection rather than the log-likelihood. Then, we use the estimates and posterior probabilities of the converged model, and perform selection of *α* and *λ* using EM. We repeat this initialization procedure for each candidate value of *K*, tuning *α* and *λ* jointly given *K* via warm starts. This procedure is described in more detail in Section 2.2.6.

#### 2.2.5. Convergence

The stopping criterion for both the CEM and EM algorithms is based upon a threshold on the relative change in the Q function. Convergence is determined when 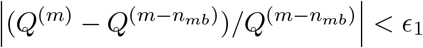 where *Q*^(*m*)^ is the value of the total Q-function conditional on current estimates at the *m^th^* EM or CEM iteration, and *n_mb_* is the minimum number of minibatches that the data can be divided into (rounded up to the nearest integer). By default, *n_mb_* = 5 (corresponding to mini-batching 20% of genes), and the left hand side (LHS) would be interpreted as the relative change in the Q-function across 5 iterations. Without mini-batching (*n_mb_* = 1), the LHS simplifies to the relative change in the Q-function across one iteration.

In FSCseq, we set *E*_1_ = 10^*−*6^ as the default convergence threshold. Convergence of IRLS and CDA are described in Section A2 and A3 of the Supplementary Material [Lim, 2020].

#### 2.2.6. Tuning the order and penalty parameters

In this section, we outline how optimal values of tuning parameters are selected in FSCseq via “warm starts” (Friedman et al., 2007; Friedman, Hastie and Tibshirani, 2010). Let (*K^*^, α^*^, λ^*^*) denote the optimal values of (*K*, *α*, *λ*), respectively. Denote candidate values of *α* as (*α*_1_*, … , α_A_*) with *α*_1_ *< · · · < α_A_*, and candidate values of *λ* as (*λ*_1_*, … , λ_L_*) with *λ*_1_ *< · · · < λ_L_*, such that there are *A · L* possible combinations of values for (*α, λ*). Also, let 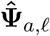 denote the final estimates of the converged results pertaining to *α* = *α_a_* and *λ* = *λ_.e_*, for a given candidate value of *K*. Then, the tuning parameter selection proceeds as follows. First, initialize a model with fixed candidate value of *K* as described in Section 2.2.4 to obtain 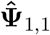. Then, leaving *α* fixed, increase *λ* to the next smallest value, *λ*_2_. Next, we run the penalized EM algorithm to convergence, utilizing the prior converged result 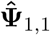 as the initial value for **Ψ**, to obtain 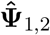. Then, we increase *λ* again and repeat the previous procedure, utilizing 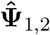 as the initial value for **Ψ**, to obtain 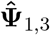. Repeat until all *L* candidate values of *λ* are searched. Then, set *α* = *α*_2_ and *λ* = *λ*_1_, and run EM to convergence from 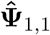 to obtain 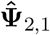. Repeat this process for each value of *α*, until all *A · L* combinations of *λ* and *α* are searched. Finally, we repeat this entire procedure for each candidate value of *K*.

For a given combination of order *K* and penalty parameters (*α*, *λ*), we may calculate the Bayesian Information Criterion (BIC) pertaining to the converged result, which has the form:

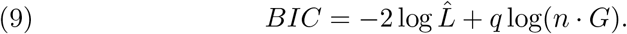

Here, 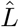 is the marginal likelihood given in (1) calculated with the estimates 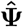 of the converged model, and *q* is the total number of estimated parameters. Let *K_j_* denote the number of distinct cluster mean parameters for gene *j*, with each fused cluster considered as one distinct cluster. Then, 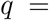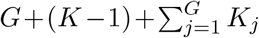, corresponding to *G* gene-level dispersion parameters, (*K −* 1) free mixing proportions (with 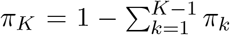, and a total of 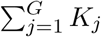 distinct log_2_ cluster means. It is clear that 1 *≤ K_j_ ≤ K* for all *j*, since *K_j_* = 1 when all clusters are fused together in gene *j* (gene is determined to be cluster-nondiscriminatory), and *K_j_* = *K* when no pair of clusters are fused together in gene *j*. Additionally, given the gene-gene independence assumption from Section 2.1, we have an effective sample size of *n·G*, corresponding to the number of entries in the gene expression matrix. To select the optimal tuning parameter values, we calculate the BIC of each converged model using (9), and select (*K^*^, α^*^, λ^*^*) as the unique combination of input values of (*K, α, λ*) corresponding to the converged model that yielded the lowest BIC.

#### 2.2.7. Prediction of discovered subtypes in new patients

FSCseq is able to perform prediction on new samples based on the converged model with input (*K^*^, α^*^, λ^*^*). Given the RNA-seq read counts of cluster-discriminatory genes in a new sample, we may derive the posterior probabilities of subtype (cluster) membership for this new sample based on (7), where we assign subtypes as done in Section 2.2.1. Specifically, we calculate the posterior probability of cluster *k* membership in this new sample, 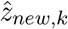, utilizing the estimated parameters from the converged FSCseq model based upon the original dataset (training set). This can be useful for predicting the subtype of new samples (prediction set) without having to re-cluster patients in future studies. We see that as in Section 2.2.1, the contribution of cluster-nondiscriminatory genes would cancel out in the posterior probability calculations of new samples.

However, correcting for sequencing depth in the new samples is nontrivial, since the size factors are specific to the training set samples used to fit the FSCseq model. To calculate the size factor of a new sample, we compare the counts of the new sample to the geometric mean of the counts of the training set samples, using the information in the training set as a pseudo-reference (Anders and Huber, 2010). Then, the estimated size factor of a new sample (*ǽ*_*new*_) is given by:

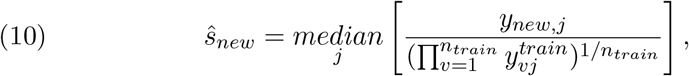

where *n_train_* denotes the training set sample size, and 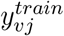 denote the count of gene *j* for the new sample and for training set sample *v*, respectively. As before, the size factors are calculated with the `DESeq2` package.

## 3. Numerical Examples

### 3.1. Simulations

To evaluate our proposed method, we simulated data across an extensive set of conditions, varying the following factors: number of true underlying clusters (*K_true_*), sample size (*n*), log_2_ fold change between cluster means (*LFC*), proportion of cluster-discriminatory genes (*p_DE_*), baseline log_2_ mean count (*β*_0_) of genes, and overdispersion (*ϕ*_0_) of counts across samples for a given gene. We fix *LFC*, *β*_0_, and *ϕ*_0_ to be the same across all simulated genes in order to demonstrate the performance of FSCseq in a controlled setting with respect to these factors. We also fixed the number of simulated genes at *G* = 10000. Of these, we specified the first (*p_DE_ · G*) genes to be cluster-discriminatory, and the rest of the genes to be nondiscriminatory, and we denote the set of cluster-discriminatory genes as *G_d_*. In cancer subtyping data, it is common to see specific genes upregulated or downregulated for just one subtype (Yersal, 2014). Thus, for each cluster-discriminatory gene *j ∈ G_d_*, we randomly select one cluster *k_*_*, 1 *≤ k_*_ ≤ K_true_*, and randomly up-regulate (*β*_*jk\**_ = *β*_0_ + *LFC*) or down-regulate (*β*_*jk\**_ = *β*_0_ *− LFC*) the log_2_ mean for that cluster, while keeping the expression of the remaining clusters the same as the baseline (*β_jk_I* = *β*_0_, for all *k^t^ /*= *k_*_*), where *β_jk_* denotes the log_2_ mean of gene *j* for a sample in cluster *k*. For each nondiscriminatory gene *j ∈/ G_d_*, we set *β_jk_* = *β*_0_ for all *k* = 1*, … , K_true_*. In this way, each cluster-discriminatory gene is expected to have one cluster with log_2_ mean different from *β*_0_, while each nondiscriminatory gene is expected to have log_2_ mean equal to *β*_0_ across all clusters.

We simulated datasets with *K_true_* = (2, 4) underlying groups with 25 and 50 samples per cluster, such that *n* = (50, 100) for *K_true_* = 2, and *n* = (100, 200) for *K_true_* = 4. Additionally, we considered *p_DE_* = (0.025, 0.050), *β*_0_ = (8, 12), *ϕ*_0_ = (0.15, 0.35, 0.50) and *LFC* = (1, 2) in our simulations. These values were determined using RNA-seq data from the TCGA Breast Cancer project (Grossman et al., 2016). Specifically, after removing outlier genes, we fit a negative binomial regression model to expression counts from each gene one-by-one, utilizing log_2_ size factors calculated from `DESeq2` as offsets and the 5 annotated PAM50 subtypes as covariates, such that a cluster mean is estimated for each subtype. The corresponding *n* × 5 design matrix (via cell-means coding) has *i^th^* row and *k^th^* column entry *x_ik_* = 1 if tumor sample *i* has annotated PAM50 subtype *k*, and *x_ik_* = 0 otherwise, for *i* = 1*, … , n* and *k* = 1*, … ,* 5. Based upon the results from these models, the 50^*th*^ and 75^*th*^ quantiles of the estimated *LFC*s in the TCGA Breast Cancer dataset were 0.875 and 1.72, and the median estimated 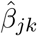 and 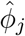 were 10.2 and 0.354, respectively. These values informed our simulation parameters above. We also simulated realistic between-sample technical variation due to sequencing depth by simulating size factors for each sample *i* = 1*, … , n* from *s_i_ ∼ N* (1, 0.25), based upon the estimates acquired by `DESeq2` on the same TCGA samples.

To prevent our simulation model from being identical to our model framework, we also introduced Gaussian noise to the log_2_ expression of each count, drawn from *σ_ij_ ∼ N* (0, 0.1) for each sample *i* and gene *j*. Then, the expression of each gene *j* = 1*, … , G* for subjects *i* = 1*, … , n* in cluster *k* = 1*, … , K_true_* was simulated from the negative binomial distribution *NB*(*μ_ijk_, ϕ*_0_) with mean *μ_ijk_*, such that

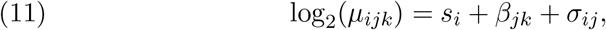

and dispersion parameter *ϕ*_0_. We simulated 100 datasets for the main simulation analyses in Section 3.1.1, and 25 datasets for the rest of the analyses.

In order to mimic what is done in real data in each simulated dataset, we used thresholds on the median count and the MAD value to pre-filter low-count and low-variable genes, respectively, keeping features that yielded median count *>* 100 and MAD score in the top 50^*th*^ quantile. We ran FSC-seq on each dataset after pre-filtering. Optimal values for tuning parameters (*K^*^, α^*^, λ^*^*) were found by searching candidate values of *K* = *{*2*, … ,* 6*}*; *α* = *{*0.01, 0.05, 0.10*, … ,* 0.50*}*; and *λ* = *{*0.25, 0.50*, … ,* 5.00*}*, as described in Section 2.2.6.

#### 3.1.1. FSCseq performance across varied conditions

In this section, we evaluate the clustering, feature selection, and prediction performance of FSCseq under a subset of simulated conditions that most closely reflected the estimates from the TCGA data. Thus, this subset represents conditions that are most similar to what one may expect from real data.

Accuracy of derived clustering results was measured by two metrics: optimal order obtained (*K^*^*) and concordance (or agreement) in cluster assignment with the truth, measured by the Adjusted Rand Index (*ARI*). Feature selection performance was measured by the true positive rate (*TPR*) and false positive rate (*FPR*) of discovering true simulated cluster-discriminatory genes. Specifically, *TPR* is the proportion of true simulated cluster-discriminatory genes (*{j*: *j ∈ G_d_}*) that were correctly determined to be discriminatory by FSCseq, while *FPR* is the proportion of true simulated nondiscriminatory genes (*{j*: *j ∈/ G_d_}*) that were incorrectly determined to be discriminatory. To assess the prediction performance of FSCseq in new data, we also independently simulated a prediction set of *n_pred_* = 25 samples for each simulated dataset, based upon the same set of simulated conditions. We then took the estimates obtained from the converged FSCseq model on the original simulated dataset (training set), and then applied the procedure from Section 2.2.7 to the corresponding test set samples to obtain the predicted cluster labels for these samples. We again measured concordance of predicted clusters with the true simulated cluster labels in the test set by *ARI*, and we denote this “prediction” *ARI*, or *pARI*. In Table 1, all performance metrics are averaged across the 100 simulated datasets, with each row denoting a unique combination of conditions. The averaged value is denoted by bar notation, e.g. 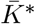 is the average *K^*^* across the 100 datasets corresponding to a particular set of simulated conditions.

**Table 1.** *Results of FSCseq clustering, feature selection, and prediction on subset of combinations of simulation conditions with K_true_* = 4 *true number of clusters and β*0 = 12 *baseline* log_2_ *mean. Clustering is measured by average optimal order 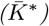 and adjusted rand index 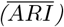. Prediction is done on an independently simulated dataset with n_pred_* = 25 *samples, and is measured by the average ARI between predicted and simulated cluster labels, and is denoted* 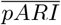. *Feature selection performance is measured by mean true positive rate* 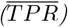 *and false positive rate* 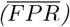 *of cluster-discriminatory gene discovery.*

| n | LFC | p<sub>DE</sub> | $\phi_0$ | $\bar{K}^*$ | $\overline{ARI}$ | $\overline{pARI}$ | $\overline{TPR}$ | $\overline{FPR}$ |
| --- | --- | --- | --- | --- | --- | --- | --- | --- |
| 100 | 1.0 | 0.025 | 0.15 | 4.05 | 0.994 | 0.998 | 0.703 | 0.0007 |
|  |  |  | 0.35 | 3.78 | 0.926 | 0.922 | 0.554 | 0.0012 |
|  |  |  | 0.50 | 2.28 | 0.233 | 0.196 | 0.077 | 0.0003 |
|  |  | 0.050 | 0.15 | 4.02 | 0.998 | 1.000 | 0.685 | 0.0006 |
|  |  |  | 0.35 | 4.01 | 0.994 | 0.994 | 0.608 | 0.0015 |
|  |  |  | 0.50 | 3.80 | 0.941 | 0.923 | 0.432 | 0.0013 |
|  | 2.0 | 0.025 | 0.15 | 4.06 | 0.996 | 0.999 | 0.780 | 0.0004 |
|  |  |  | 0.35 | 4.01 | 0.999 | 1.000 | 0.757 | 0.0004 |
|  |  |  | 0.50 | 4.02 | 0.998 | 1.000 | 0.726 | 0.0014 |
|  |  | 0.050 | 0.15 | 4.05 | 0.995 | 0.999 | 0.781 | 0.0012 |
|  |  |  | 0.35 | 4.04 | 0.997 | 0.999 | 0.747 | 0.0010 |
|  |  |  | 0.50 | 4.01 | 0.999 | 1.000 | 0.725 | 0.0026 |
| 200 | 1.0 | 0.025 | 0.15 | 4.17 | 0.988 | 0.993 | 0.741 | 0.0014 |
|  |  |  | 0.35 | 4.07 | 0.989 | 0.992 | 0.700 | 0.0018 |
|  |  |  | 0.50 | 3.97 | 0.975 | 0.974 | 0.645 | 0.0012 |
|  |  | 0.050 | 0.15 | 4.15 | 0.992 | 0.996 | 0.727 | 0.0017 |
|  |  |  | 0.35 | 4.04 | 0.997 | 0.997 | 0.700 | 0.0034 |
|  |  |  | 0.50 | 4.00 | 1.000 | 1.000 | 0.662 | 0.0014 |
|  | 2.0 | 0.025 | 0.15 | 4.11 | 0.992 | 0.999 | 0.803 | 0.0019 |
|  |  |  | 0.35 | 4.07 | 0.998 | 1.000 | 0.798 | 0.0004 |
|  |  |  | 0.50 | 4.09 | 0.995 | 0.999 | 0.782 | 0.0007 |
|  |  | 0.050 | 0.15 | 4.05 | 0.996 | 0.999 | 0.809 | 0.0019 |
|  |  |  | 0.35 | 4.04 | 0.997 | 0.999 | 0.778 | 0.0007 |
|  |  |  | 0.50 | 4.04 | 0.998 | 0.998 | 0.767 | 0.0005 |

We found that the clustering performance of FSCseq was generally robust to the magnitude of *ϕ*_0_, even when *n* is small (Table 1). One exception was found when *n* = 100, *p_DE_* = 0.025 and *LFC* = 1 with *K_true_* = 4, i.e. when both the proportion of discriminatory genes and the *LFC* across these discriminatory genes were small. In such a case, the clusters may be poorly separated and clustering results may be confounded by higher *ϕ*_0_, compounded by the small average per-cluster sample size (25 per cluster). We note that none of the compared methods perform well in this case (see Section C2 of the Supplementary Material [Lim, 2020]), although FSCseq yields higher mean *ARI*.

As *n*, *LFC*, or *p_DE_* is increased, we also found that that the sample clusters become more distinct from one another, yielding better clustering results. Increasing *n* and *LFC* also tended to result in an increase in *TPR*, reflecting the increasing sensitvity of our variable selection method under larger sample size and fold change differences between clusters. However as *ϕ*_0_ increased, we found a decrease in the *TPR*, where the higher overdispersion confounded the true differences in expression of the cluster-discriminatory genes and reduced our sensitivity to select cluster discriminatory genes. Interestingly, the *FPR* did not vary significantly across changes in these conditions, showing the reliability of cluster-discriminatory features discovered by FSCseq, and the *FPR* was relatively low in general. Figure 1 shows a scatterplot of *TPR* vs *FPR* from each individual simulated dataset from Table 1, stratified by simulated *LFC* and *n*. In addition to showing that larger *n* or *LFC* and a smaller value of *ϕ*_0_ yielded higher *TPR*, we also show that selecting the correct order was important in attaining both high *TPR* and low *FPR*. This is because FSCseq more accurately selected cluster-discriminatory genes if the correct number of groups is identified.

**Fig 1.**
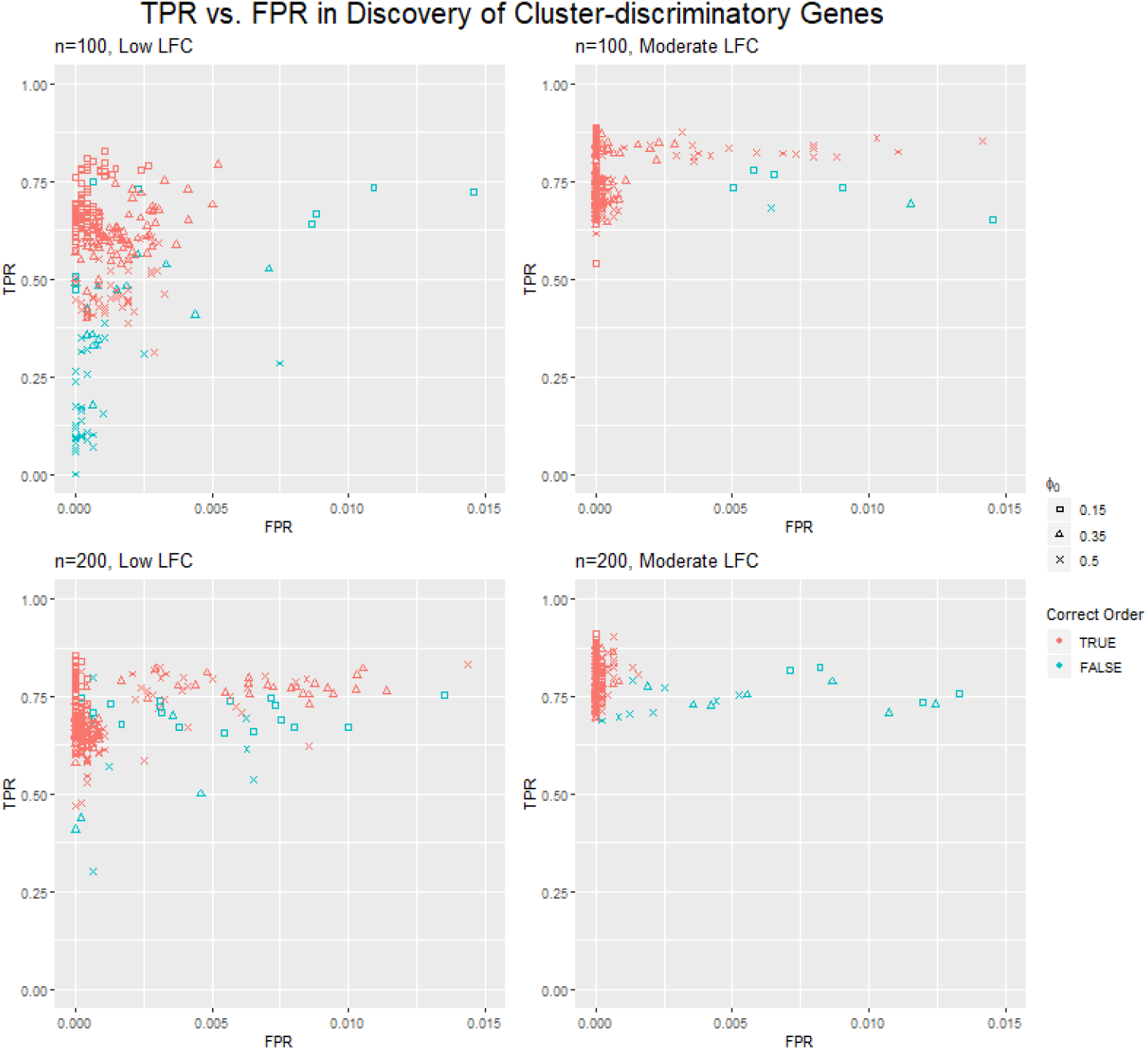
*Scatterplot of true positive rate (TPR) vs false positive rate (FPR) in discovering cluster-discriminatory genes in simulated datasets via FSCseq. Displayed points correspond to simulated datasets with K_true_* = 4 *underlying clusters, with n* = 100 *(top) or* 200 *(bottom) and simulated LFC* = 1 *(‘Low’, left) or* 2 *(‘Moderate’, right). Red points indicate that the correct order was uncovered (K^*^* = *K_true_), and blue points indicate that an incorrect order was uncovered. Squares, circles, and X’s indicate low (ϕ*_0_ = 0.15*), moderate (ϕ*_0_ = 0.35*), and high (ϕ*_0_ = 0.50*) levels of overdispersion, respectively. In general, TPR is higher for larger n and larger LFC. TPR is higher and FPR is lower for results that yielded the correct order, or for smaller values of ϕ*_0_*. For ease of visualization, a total of 15 outlier points (of 2400 total) were removed in this figure with* 0.015 *< FPR <* 0.060*. The figure without excluding outlier points can be found in Supplementary Figure 2, Section B of the Supplementary Material [Lim, 2020].*

FSCseq’s prediction performance was also found to be highly dependent on its initial clustering performance during model training, i.e., *pARI* on simulated test subjects was correlated to *ARI* during FSCseq training. This reflects the fact that FSCseq models that correctly clustered samples also tended to yield more accurate parameter estimates, which allowed for more accurate prediction of cluster identity in new samples. Since FSCseq generally yielded high *ARI*, we also generally found high average *pARI* for simulated conditions, reflecting the accuracy of predictions via FSCseq. Overall, we found favorable performance of FSCseq over a variable series of conditions.

Results of FSCseq analyses on all simulated datasets encompassing all simulation conditions can be found in Section B of the Supplementary Material [Lim, 2020].

#### 3.1.2. Performance comparison across methods

We compared performance of FSCseq with 7 competing methods: `iCluster+` (iCl), average-linkage hierarchical clustering (HC), K-medoids (KM), `NB.MClust` (NBMB), and mclust on log, variance stabilizing, and rlog transformations of the normalized data (lMC, vMC, and rMC). FSCseq can intrinsically adjust for differences in sequencing depth using derived size factors as described earlier. For all other methods, we use either normalized counts (iCl, HC, KM, NBMB, lMC) or transformed normalized counts (vMC, rMC) for clustering to adjust for differences in sequencing depth, and these counts are computed from `DESeq2` (Love, Huber and Anders, 2014). We also note that KM was performed on log-transformed normalized counts (Jaskowiak, Costa and Campello, 2018) as in lMC. Because there was no default method of selecting *K^*^* for HC, we chose to utilize the gap statistic, as proposed by Tibshirani, Walther and Hastie (2001), and implemented the order selection in this setting via the `NbClust` R package (Charrad et al., 2014). For KM, we selected *K^*^* to be the value of *K* that maximized the average silhouette width (Reynolds, Richards and Rayward-Smith, 2004). For NBMB and the transformed MC methods, we used default selection procedures for *K^*^* directly available from the respective `NB.MClust` and mclust R packages. For `iCluster+`, we describe an automated procedure for selecting *K^*^* in Section C1 of the Supplementary Material [Lim, 2020].

Cluster-discriminatory gene selection performance was not compared across methods as only FSCseq had an automated procedure for feature (gene) selection. We note that `iCluster+` does perform automatic gene selection via the L1 penalty; however, the `iCluster+` manual recommends thresholding an arbitrary proportion of genes for selection rather than utilizing its automatic feature selection process (Mo and Shen, 2019). One could similarly perform thresholding on the estimated coefficients from the mclust or `NB.MClust` results, but this would also depend on an arbitrary threshold. We compared the clustering performance of these methods under different numbers of underlying groups *K_true_* = (2, 4) with 25 samples per cluster. We also varied *ϕ*_0_ to test each method’s robustness to the level of overdispersion in the simulated counts. Here, we define order accuracy (*OA*) as the proportion of simulated datasets (within a given simulated condition) that yielded the correct order, such that *K^*^* = *K_true_*.

Figure 2 shows violin plots of order accuracy *OA* and *ARI*. Here, we fixed *LFC* = 1, *p_DE_* = 0.05, *n/K_true_* = 25, and *β*_0_ = 12, and we varied *ϕ*_0_ = *{*0.15, 0.35, 0.50*}*. Generally, FSCseq yielded clusters that had the highest concordance with the simulated clusters. Many of the methods performed competitively in clustering when *K_true_* = 2, but FSCseq and `iCluster+` performed markedly better than competing methods when *K_true_* = 4. Throughout all conditions, `iCluster+` tended to select a larger optimal order *K^*^ > K_true_*, which caused performance to be significantly better under *K_true_* = 4 than *K_true_* = 2. KM performed sporadically throughout, suggesting lack of robustness in clustering performance to the magnitude of *ϕ*_0_. Generally, NBMB yielded the lowest *ARI* compared to all other methods. We postulate that this is because NBMB utilizes cluster-specific dispersion parameters for each gene in their model, which may yield unstable estimates due to smaller sample sizes in estimating each cluster-level dispersion parameter.

**Fig 2.**
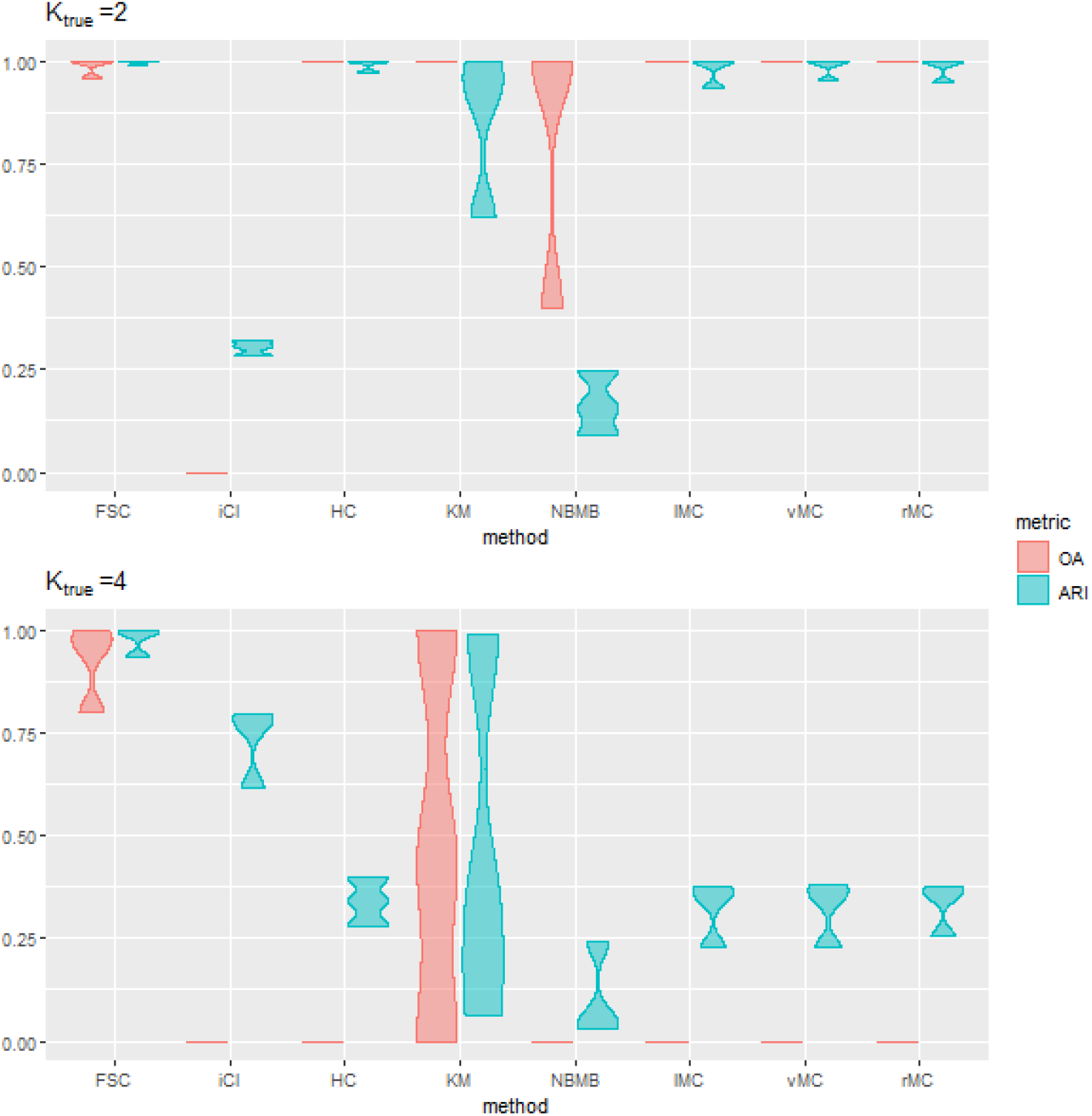
*Violin plots of Order Accuracy (OA, in red) and average cluster concordance with the truth (ARI, in blue) from simulation results with K_true_* = 2 *and K_true_ = 4. We compared performance of FSCseq (FSC), `iCluster+` (iCl), hierarchical clustering (HC), K-medoids (KM), `NB.MClust` (NBMB), and mclust on log/variance-stabilizing/rlog transformed data (lMC/vMC/rMC). Simulated overdispersion was varied with ϕ*_0_ = (0.15, 0.35, 0.50) *to test each method’s robustness to the magnitude of overdispersion. Other simulation parameters were fixed at LFC* = 1*, p_DE_* = 0.05*, β*0 = 12*, and n/K_true_* = 25 *for both K_true_* = 2 *(top) and K_true_* = 4 *(bottom). We fix n/K_true_ here to show the effect of varying K_true_ on performance, with a fixed number of samples per cluster. Many methods perform competitively when K_true_* = 2*, but FSCseq attains the highest average ARI overall, and yields high performance that is very robust to the magnitude of ϕ*_0_.

In general, we observed poorer clustering performance of most methods in *K_true_* = 4 compared to *K_true_* = 2, with markedly lower average *ARI* for HC and the transformed MC runs (Figure 2). When *K_true_* is larger, there is a smaller proportion of samples within each cluster-discriminatory gene with log_2_ mean different from *β*_0_, since only one cluster is differentially regulated by *LFC* for these genes. Thus, we generally observe a significant decrease in performance for larger *K_true_*, although FSCseq’s performance is most robust to this effect. In addition, similarly sporadic performance of HC in RNA-seq was also observed previously (Vidman, Källberg and Rydén, 2019), suggesting unreliability of HC for clustering RNA-seq gene expression. Finally, although the MC methods performed similarly on average, the best performing method between these three transformations varied for each set of simulation conditions, reflecting the fact that the optimal transformation may not be known in advance. Comparisons between FSCseq and these competing methods across the entire extensive set of simulation conditions can be found in Section C2 of the Supplementary Material [Lim, 2020].

#### 3.1.3. Performance under simulated batch effects

We also evaluated the clustering and order selection performance of these methods in the presence of batch effects. Here we simulate two batches in a subset of simulation conditions, where batch effects were imposed on a subset of genes for each simulated dataset. Simulated RNA-seq read counts were generated in a manner similar to (11), except now we assume log_2_(*μ_ijk_*) = *s_i_* +*β_jk_* +*σ_ij_* +*{I*(*j ∈ G_b_*)*· γ*_0_ *· Batch_i_}*, where *G_b_* is the set of genes that are simulated to be batch-affected, *γ*_0_ is a fixed batch effect on the log_2_ scale, and *Batch_i_* = *{−*0.5, 0.5*}*. We randomly assign *Batch_i_* = *−*0.5 or *Batch_i_* = 0.5 for *i* = 1*, … n* with equal probability, and *G_b_* is comprised of a randomly selected 50% subset of the *G* genes. For each simulation we performed FSCseq with the batch variable as a covariate, and compared performance via each method on simulation conditions with *n/K_true_* = 50, *LFC* = 2, *p_DE_* = 0.05, *β*_0_ = 12, and *ϕ*_0_ = 0.35. We chose this subset to isolate the confounding effect of batch, since these simulated conditions yielded good clustering performance when batch effects were not simulated.

Results on batch-simulated datasets are shown in Table 2. FSCseq is the only method that performs robustly to the magnitude of *γ*_0_, showing its ability to properly adjust for the confounding batch effects. Although KM does not perform any correction for batch, it surprisingly performed well under *K_true_* = 2 and *γ*_0_ = 2. However, because KM does not actually correct for batch, KM performs expectedly poorly under larger simulated batch effects *γ*_0_ = 3.

**Table 2.** **Average obtained order** 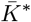 *and average ARI for competing methods with K_true_* = 2 *and K_true_* = 4 *underlying groups, in the presence of simulated batch effects γ*0*. We shorten notation in this table by denoting the 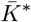 value as 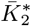 for K_true_* = 2*, and as 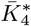 for K_true_* = 4*. Similarly, we denote ARI as ARI*2 *for K_true_* = 2*, and as ARI*4 *for K_true_* = 4*. Tabulated results are from datasets with n/K_true_* = 50 *simulated samples per cluster, with fixed LFC* = 2*, pDE* = 0.05*, and β*0 = 12*, ϕ*_0_ = 0.35*. Simulated effect across batches was varied, such that γ*0 = 2, 3.

| | $\gamma_0$ | $\bar{K}_2^*$ | $\overline{ARI}_2$ | $\bar{K}_4^*$ | $\overline{ARI}_4$ |
| --- | --- | --- | --- | --- | --- |
| FSC | 2 | 2.00 | 1.000 | 4.16 | 0.971 |
|  | 3 | 2.00 | 1.000 | 4.00 | 0.965 |
| iCl | 2 | 4.68 | 0.450 | 5.00 | 0.779 |
|  | 3 | 3.00 | 0.287 | 4.00 | 0.332 |
| HC | 2 | 2.00 | 0.000 | 2.00 | 0.001 |
|  | 3 | 2.00 | -0.002 | 2.00 | -0.001 |
| KM | 2 | 2.00 | 0.989 | 2.00 | 0.002 |
|  | 3 | 2.00 | -0.002 | 2.00 | -0.001 |
| NBMB | 2 | 2.28 | 0.055 | 2.32 | 0.004 |
|  | 3 | 2.32 | 0.043 | 2.04 | -0.001 |
| lMC | 2 | 3.92 | 0.498 | 2.00 | 0.001 |
|  | 3 | 4.00 | 0.499 | 2.00 | -0.001 |
| vMC | 2 | 3.92 | 0.498 | 2.00 | 0.001 |
|  | 3 | 4.00 | 0.499 | 2.00 | -0.001 |
| rMC | 2 | 2.04 | 0.024 | 2.00 | 0.001 |
|  | 3 | 3.36 | 0.368 | 2.00 | -0.001 |

### 3.2. Application to TCGA Breast Cancer RNA-seq dataset

In this section, we compared performance of these clustering methods on the TCGA Breast Cancer (BRCA) dataset. The dataset and annotations were obtained from the National Cancer Institute GDC Portal (Grossman et al., 2016). Gene expression RNA-seq data was sequenced via the Illumina Hiseq 2000 platform, and alignment and quantification were done using the GDC’s mRNA Analysis Pipeline, which can be found in the “GDC Data User’s Guide” online (Grossman et al., 2016). The raw read counts were downloaded using the `TCGAbiolinks` R package (Colaprico et al., 2015).

Subtyping was done previously using the PAM50 classifier: a set of 50 genes that has been heavily investigated as driving genes of breast cancer subtypes (Koboldt et al., 2012). These genes are also known to be primarily expressed in tumor cells. However, tumor bulk samples are typically comprised of heterogeneous mixtures of cell types, and may poorly reflect the expression profile of the tumor if the sample is of low tumor purity, i.e. if it is composed of a small proportion of tumor cells. Attempts at unsupervised clustering of low-purity samples based on subtype may emphasize genes that are associated with purity and confound results (Aran, Sirota and Butte, 2015). Thus, we first included in our analysis only samples whose estimated ABSOLUTE purity (Carter et al., 2012) was greater than 0.9. This is done in order to (1) properly cluster samples via each compared method based upon tumor subtypes, and (2) compare each clustering method fairly with the annotated subtypes, which are based upon tumor-intrinsic PAM50 classifier genes. We also performed our analyses on all samples without purity filtering, and compared our results with those from Koboldt et al. (2012), which performed similar clustering analyses on all samples.

After filtering samples by purity, we found very low incidence of the HER2-enriched and normal-like subtypes, and thus removed these subtypes from our analysis. Then, we pre-filtered low read count genes by low median normalized count, and low-variable genes by low MAD scores. For reproducibility, we detail the pre-processing steps in Section D1 of the Supplementary Material [Lim, 2020]. After all filtering steps, the dataset contained 123 samples from 3 distinct subtypes, and 4038 genes. We measured concordance with annotated sample groupings (tumor subtypes in the TCGA BRCA dataset) by four clustering metrics: ARI, normalized mutual information (NMI) (Strehl and Ghosh, 2003), normalized variation of information (NVI) (Reichart and Rappoport, 2009), and normalized information distance (NID) (Vinh, Epps and Bailey, 2010).

The RNA-seq experiments were performed on separate wells of multiwell plates, with multiple samples sequenced on the same plate at different times. Therefore, samples across plates may express variability similar to batch effects (Reese et al., 2013). We performed FSCseq analysis with (*FSC_adj_*) and without (*FSC*) adjustment for this confounding effect, and compared the results with competing methods. For proper analysis via *FSC_adj_*, we agglomerated singleton plates (plates with just one sample) into one joint plate.

For this data, we expanded the grid of candidate values of *K* to *{*2*, … ,* 8*}*. All clustering metrics were calculated with respect to the mRNA PAM50 subtype annotations (‘*anno*’) by the GDC. Potential confounders like age, ethnicity, and tumor grade were not included here because they were not of major interest in our analysis, but they can be incorporated as additional covariates in FSCseq. Results are given in Table 3.

**Table 3.**
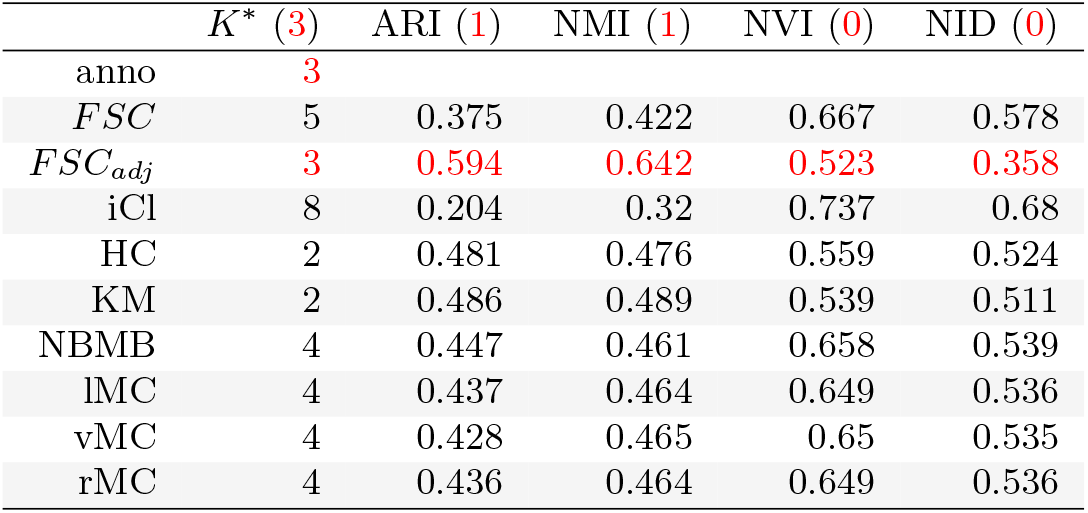
Selected order (K^*^) and clustering concordance between compared methods and annotated TCGA Breast Cancer subtypes. FSCseq was run with adjustment (FSC_adj_) and without adjustment (FSC) for plate effect, and each of the clustering labels were compared to annotated subtypes (anno). For each column, the value of the best performing metric is colored in red. The values in parentheses in the column headings represent the optimal value for that metric. For ARI and NMI, values close to 1 indicate good clustering, and values close to 0 indicate poor clustering. For NVI and NID, values close to 0 indicate good clustering, and values close to 1 indicate poor clustering.

*FSC* selected order *K^*^* = 5, but after adjusting for plate effects, *FSC_adj_* uncovered the true order of *K^*^* = 3. Similar to the trends seen in simulations, `iCluster+` selected a larger number of clusters *K^*^* = 8, while HC and KM selected a smaller *K^*^* = 2. The transformed MC methods and NBMB selected a slightly larger *K^*^* = 4. Of the competing methods, *FSC_adj_* yielded the clusters with the highest concordance with the annotations, with an *ARI* of 0.594, followed by KM and HC with *ARI*s of 0.486 and 0.481, respectively. *FSC_adj_* additionally yielded the best *NMI*, *NV I*, and *NID*.

Feature selection via *FSC* and *FSC_adj_* respectively determined a total of 2693 and 2238 genes to be cluster-discriminatory, while the automatic feature selection by the L1 penalty in `iCluster+` determined significantly more genes (4035) to be cluster-discriminatory. Of the 41 PAM50 genes that were included in our analysis after pre-filtering, *FSC* determined all 41 likely to be cluster-discriminatory, and *FSC_adj_* determined 38 of them to be likely. This shows FSCseq’s ability to identify subtype-discriminating genes in real cancer data, as the relevance of PAM50 genes in RNA-seq has been shown by previous studies (Picornell et al., 2019; Raj-Kumar et al., 2019). Figure 3 shows a heatmap of the PAM50 genes with notations on their inclusion/exclusion through the pre-filtering step, and through FSC-seq’s simultaneous feature selection. Column (samples) ordering is based on annotated subtypes, and samples are ordered within subtypes by decreasing order of maximum posterior probability from *FSC_adj_* results. KM performs well in terms of ARI, but it only grouped samples by basal vs. non-basal subtypes and didn’t distinguish between the Luminal A and Luminal B subtypes. Similarly, lMC performs well in grouping basal samples, but over-selected the order (*K^*^* = 4) and was not able to distinguish between Luminal A and B samples. Overall, the cluster labels from *FSC_adj_* most accurately clustered the samples according to the 3 underlying subtypes. Additional results from all cluster-discriminatory genes with all compared methods can be found in Section D2 of the Supplementary Material [Lim, 2020].

**Fig 3.**
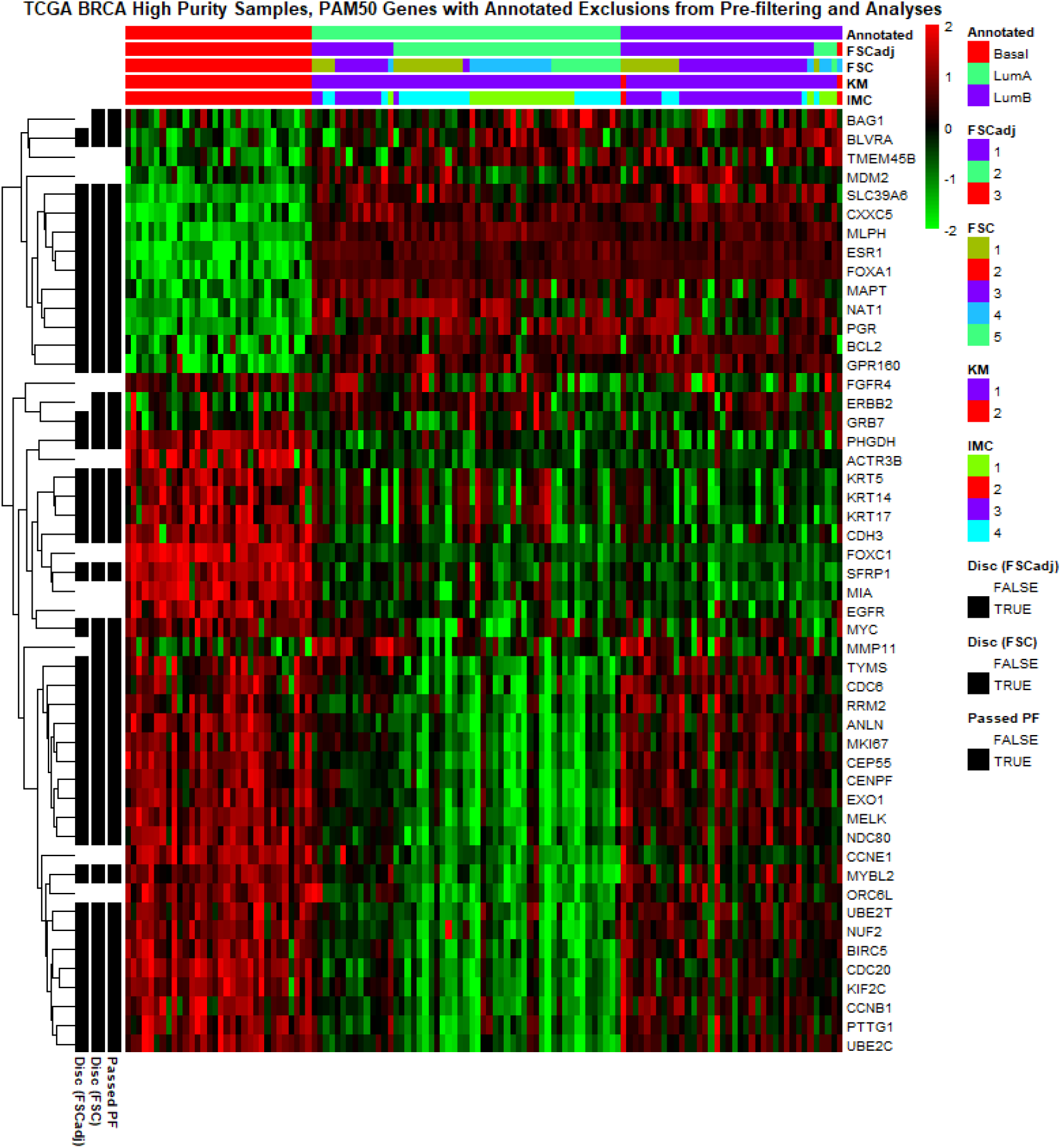
Heatmap of the PAM50 genes included in FSCseq analyses, with row annotations for feature selection and pre-filtering (left) and column annotations for clustering labels (top). Column ordering is based on annotated subtypes, and samples are ordered within subtypes by decreasing order of maximum posterior probability from FSC_adj_ results. 9 PAM50 genes did not pass the pre-filtering (PF) threshold: TMEM45B, MDM2, FGFR4, ACTR3B, FOXC1, MIA, EGFR, CCNE1, and ORC6L. Of the 41 remaining PAM50 genes, FSC and FSC_adj_ found 41 and 38 of the PAM50 genes were cluster-discriminatory, respectively. Additionally, all clustering labels distinguish well between Basal and Luminal subtypes, but FSC_adj_ best distinguishes between Luminal A and Luminal B samples.

In addition to the PAM50 genes, other significant cluster-discriminatory genes found by both *FSC* and *FSC_adj_* included genes like CDH1, which has been linked to lobular or ER+ tumor carcinomas (Yang, Ding and Huang, 2015), and CHEK2, which has been studied for its association with the luminal subtypes (Huszno and Kolosza, 2019). Additionally, we performed gene ontology analysis on the discovered sets of cluster-discriminatory genes from both *FSC* and *FSC_adj_*. Enriched pathway analysis showed that genes from both sets were involved in known key pathways associated with cancer. Such pathways included signaling pathways like EIF2 and AHR, as well as pathways involved in mitosis and other molecular mechanisms of cancer. Both sets of genes also contained many gene ontology (GO) biological processes pertaining to the cell cycle, consistent with previous studies that found such enrichment in basal-like subtypes of breast cancer (Yang et al., 2017; Yang, Gao and Luo, 2019). We further validated our results by testing for overlaps with known gene sets via GSEA analysis (Mootha et al., 2003; Subramanian et al., 2005). We first grouped cluster-discriminatory genes from *FSC_adj_* using *MBCluster.Seq* (Si et al., 2013), then separately analyzed the subset of genes that were upregulated (*basalUP*), and those that were downregulated (*basalDOWN*) for the most distinct basal subtype. Significantly overlapping gene sets with *basalUP* and *basalDOWN* were sets of genes known to differentiate subtypes of breast cancer. Additionally, the top overlapping gene set for *basalUP* and *basalDOWN* corresponded to the specific collections of genes known to be upregulated and downregulated, respectively, in the basal subtype of breast cancer (Smid et al., 2008). Detailed results of our gene enrichment analyses can be found in Section D3 of the Supplementary Material [Lim, 2020].

We measured performance in prediction via leave-one-out cross-validation by training a model on cluster-discriminatory genes from *FSC* with one sample held out, and using this model to predict the cluster label of the held-out sample. In this way, prediction was performed on each sample once. We observed very high overall accuracy of 0.951, showing robustness of our prediction framework in real data settings.

Finally, we also clustered the TCGA samples without pre-filtering samples for purity and including all 5 original subtypes (*K_true_* = 5), as done previously by Koboldt et al. (2012). We anticipated that there would exist a larger number of underlying groups in this dataset due to heterogeneity caused by low purity samples, which is also reflected by the results in Koboldt et al. (2012). Thus, we expanded the search range of *K* to *K* = *{*2*, … ,* 15*}*. As expected, all methods yielded clusters of poor concordance with the annotated PAM50 subtypes, with *FSC* and *FSC_adj_* yielding larger orders of *K^*^* = 7 and *K^*^* = 8, respectively, and *ARI*s of 0.316 and 0.245, respectively. Koboldt et al. (2012) performed unsupervised clustering on the same samples and found 13 clusters, and they performed semi-supervised clustering with an intrinsic list of significant genes that similarly yielded 14 clusters. Their results were similarly poor in agreement with the annotated subtypes (*ARI* of 0.272 and 0.258 for unsupervised and semi-supervised clusters, respectively). Compared to these results, FSCseq was able to select *K^*^* that is closer to the true number of subtypes, however the potential confounding in expression due to variable sample purity appears to cloud the ARI performance of all methods in this setting. All results from this analysis can be found in Section D4 of the Supplementary Material [Lim, 2020].

## 4. Discussion

Our findings from our simulations and real data applications give evidence to the utility of our method across a varied set of conditions. In our simulations, we found very good feature selection performance throughout simulated conditions, and FSCseq outperformed existing clustering methods for RNA-seq data. In addition, the markedly low *FPR*s in gene discovery may help researchers attain a higher degree of confidence in the discovered list of genes, while limiting expended resources on validation studies in clinical settings. In the TCGA BRCA dataset, FSCseq clusters aligned best to previously discovered subtypes when correcting for batch effects (plate), although differing levels of heterogeneity in samples confounded the results. The annotated subtypes were discovered by analyzing just the genes that were clinically validated to be significant in discriminating across breast cancer subtypes. We showed that FSCseq is able to perform comparably, despite using no such *a priori* knowledge of significant genes. Moreover, of the PAM50 genes included in our TCGA BRCA analysis, the FSCseq workflow identified most of these genes as discriminatory across the resulting clusters. As these genes have been validated to be significant on both microarray and RNA-seq platforms, this emphasizes FSCseq’s ability to uncover significant genes in real data settings.

## SUPPLEMENTARY MATERIAL

**Supplement A: Supporting Information for Model-based Feature Selection and Clustering of RNA-seq Data for Unsupervised Subtype Discovery** (doi: COMPLETED BY THE TYPESETTER;.pdf). Contains links to code and implementation of FSCseq and additional details of our algorithm in Section A, extensive results from simulations in Sections B-C, and details and results from the TCGA BRCA dataset in Section D.

