## Supplementary Materials for "Model-Based Feature Selection and Clustering of Rna-Seq Data for Unsupervised Subtype Discovery"

#### Section A: Additional Details of FSCseq

The workflow for FSCseq and an example visualization of the fusion SCAD penalty are given in Figure 1. The `FSCseq` R package for implementation of our method can be found here: https://github.com/DavidKLim/FSCseq/. R code to perform all analyses can be found here: https://github.com/DavidKLim/FSCseqPaper/. All resources necessary to replicate our results can be obtained by following the `FSCseqPaper` code. The TCGA BRCA dataset used in our analysis was downloaded on March 25th, 2019.

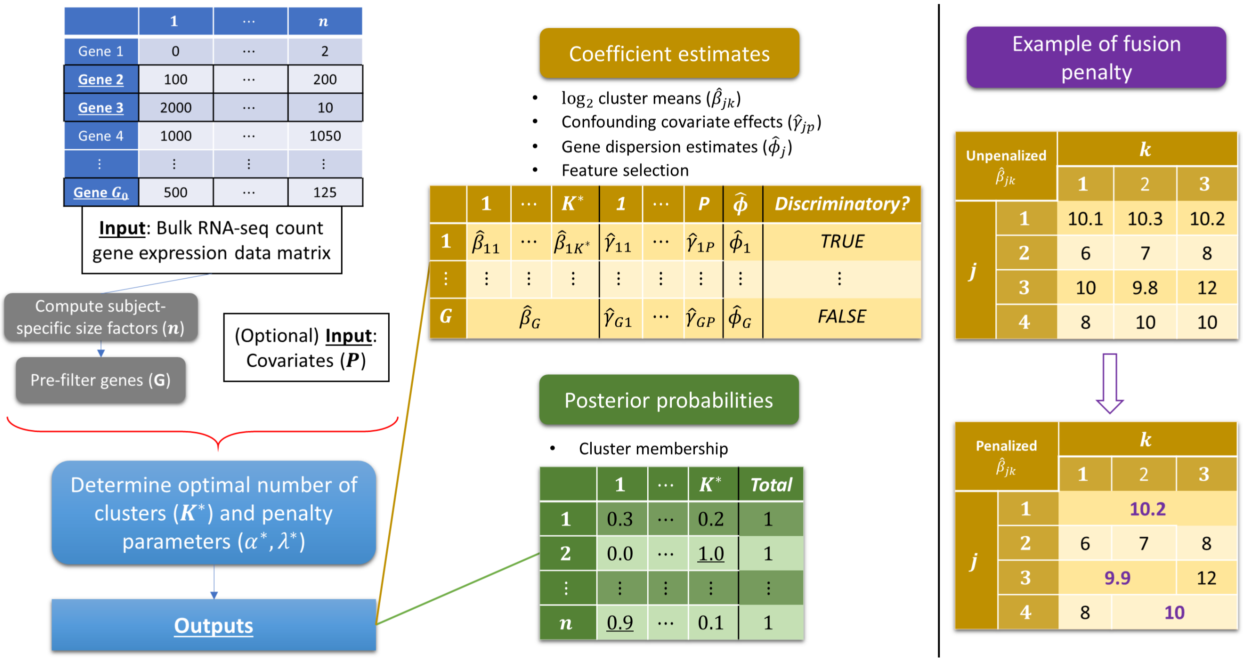


Supplementary Figure 1: Example workflow of FSCseq (left) and visualization of the mechanics of the fusion SCAD penalty (right) imposed on $\hat{\beta}_{jk}$ with $j = 1, \ldots, 4$ and $k = 1, \ldots, 3$. For prediction, the coefficient estimates and the list of cluster-discriminatory genes obtained from FSCseq can be used to compute posterior probabilities of cluster membership on new samples. Importantly, the fusion penalty is imposed on each gene separately, and clusters are fused together within each given gene separately. When all clusters are fused together for a particular gene ($j = 1$, right), that gene is determined to be nondiscriminatory across clusters. Otherwise, the gene is considered cluster-discriminatory.

#### A1: SCAD Fusion Penalty

The SCAD fusion penalty given in Section 2.1.1 is dependent on the quadratic penalty method (Nocedal and Wright, 2000) and the SCAD penalty (Fan and Li, 2001). The magnitudes of the two penalty terms are controlled by tuning parameters $\lambda$ and $\alpha$. Intuitively, $\lambda \geq 0$ controls the overall amount of the penalty incorporated in the model, while $\alpha \in (0, 1)$ balances between the quadratic penalty term ($\alpha = 0$) and the SCAD penalty term ($\alpha = 1$).

The form of the SCAD penalty is most commonly given by its first derivative: $SCAD'_{\lambda^*}(\theta) = \lambda^*\{I(\theta \leq \lambda^*) + [(a\lambda^* - \theta)_+ / (a\lambda^* - \lambda^*)] I(\theta > \lambda^*)\}$ for $a > 2$, $\theta > 0$, and $\lambda^* = \lambda\alpha$. As suggested by Fan and Li (2001), we set $a = 3.7$.

Let $k_*$ denote a dummy index for the cluster consisting of data from clusters $k$ and $l$. When $\hat{\theta}_{j,kl} = 0$, then $\hat{\beta}_{jk_*}$ is estimated based upon the data in both clusters $k$ and $l$, and is set equal to the $\log_2$ means for clusters $k$ and $l$ for that gene $j$: $\hat{\beta}_{jk_*} = \hat{\beta}_{jk} = \hat{\beta}_{jl}$, thus setting $\hat{\beta}_{jk} = \hat{\beta}_{jl}$ when $\hat{\theta}_{j,kl} = 0$. We do this by adding the posterior probabilities for each sample in the fused clusters for the current EM/CEM iteration. Then, we proceed with the EM/CEM algorithm as before, maximizing relevant parameters for the fused cluster $k_*$ in gene $j$ using the data from both clusters $k$ and $l$. The procedure is generalized in the same fashion when more than 2 clusters are fused. It is important to note that we do not fuse two clusters together for every gene, but fuse clusters independently within each gene, depending on the estimates of the $\log_2$ means across clusters for that gene. We use the BIC to tune $\alpha$ and $\lambda$ (details in Section 2.2.6 of the main text) to search for the optimal magnitude of the quadratic penalty constraint. We show that this method works well in simulations and real data.

#### A2: Iteratively re-weighted least squares

The method of iteratively reweighted least squares (IRLS) uses transformed responses that are reweighted across each iteration of IRLS (Breheny and Huang, 2011). For fixed gene $j$, we transform the responses by

$$\tilde{y}_{ijk} = \eta_{ijk} + (\frac{y_{ij} - g(\eta_{ijk})}{g'(\eta_{ijk})}), \tag{1}$$

where $g(\cdot)$ is the inverse link function for the Negative Binomial family with $\log_2$ link, thus $g(\eta_{ijk}) = \mu_{ijk}$, where $g^{-1}(\mu_{ijk}) = log_2(\mu_{ijk})$ is the $\log_2$ link function given in Equation 2 under Section 2.1 of the main text.

Let $\mathbf{\Theta}_j$ be a vector of length $(K+P)$ of the $K$ cluster $\log_2$ means and the $P$ covariates for a fixed gene $j$: $\mathbf{\Theta}_j = (\beta_{j1}, \ldots, \beta_{jK}, \gamma_{j1}, \ldots, \gamma_{jP})^T$. Incorporating the above transformation and taking the Taylor series expansion of the coefficient estimates around current estimates of $\mathbf{\Theta}_j$, the portion of the Q-function pertaining to the $j$'th gene becomes the typical IRLS form for generalized linear models (McCullagh and Nelder, 1989):

$$Q_j(\mathbf{\Theta}_j) \approx \frac{1}{2n}(\tilde{\mathbf{y}}_j - \mathbf{X}\mathbf{\Theta}_j)'\mathbf{W}_j(\tilde{\mathbf{y}}_j - \mathbf{X}\mathbf{\Theta}_j) + p_{\lambda,\alpha}(\boldsymbol{\beta}_j), \tag{2}$$

where $\tilde{\mathbf{y}}_j$ is a vector $(\tilde{y}_{1j1}, \ldots, \tilde{y}_{nj1}, \tilde{y}_{1j2}, \ldots, \tilde{y}_{nj2}, \ldots\ldots, \tilde{y}_{njK})$ of length $nK$ for a fixed gene $j$, corresponding to the $n$ observations repeated once for each of the $K$ clusters. $\mathbf{X}$ is a matrix of dimensions $(nK) \times (K+P)$, where the first $K$ columns represent the design matrix for the intercept-only model using cell means coding with each cell representing a cluster, and with $n$ replicates for each cell. In particular, columns of X for $k = (1, \ldots, K)$ is given by: $x_{ik} = I(i \in [(k-1)n+1, kn])$. The last $P$ columns represent the design matrix of covariate values for each sample, where each sample's measurement is repeated $K$ times, once for each cluster. $\mathbf{W}_j$ is a $nK \times nK$ diagonal matrix with the IRLS weights on the diagonal to correspond to each element of the transformed response, where the diagonal of $\mathbf{W}_j$ are the corresponding IRLS weights, given by: $w_{ijk} = \sqrt{\frac{\hat{z}_{ik}^{(m)} g'(\eta_{ijk})^2}{V(\mu_{ijk})}}$, and the variance is $V(\mu) = \mu + \phi\mu^2$ for the Negative Binomial. Then, the corresponding diagonals of $\mathbf{W}_j$ are $(w_{1j1}, \ldots, w_{nj1}, w_{1j2}, \ldots, w_{nj2}, \ldots\ldots, w_{njK})$. The form of the fusion SCAD penalty $p_{\lambda,\alpha}(\boldsymbol{\beta}_j)$ can be found in Equation 5 of the main text.

We note in particular that the first term of Equation 2 is quadratic, and thus convex and differentiable. Thus, the $\theta_{j,kl} = \beta_{jk} - \beta_{jl}$ reparametrization of our fusion penalty allows for convexity of the overall gene-specific penalized Q function, guaranteeing convergence to a global maximum via coordinate descent (Tseng, 2001). As explained in the main text, we maximize $\hat{\boldsymbol{\beta}}_{\boldsymbol{j}}$ and $\hat{\boldsymbol{\gamma}}_{\boldsymbol{j}}$, using $\hat{\boldsymbol{\theta}}_{\boldsymbol{j}}$ to facilitate the coordinate-descent algorithm (CDA), which is nested within the IRLS algorithm. In the CDA (inner) loop, we sequentially update each parameter while fixing all others at their current iteration's estimates In the IRLS (outer) loop, we recompute the IRLS weights matrix $\mathbf{W}_j$ and compute (2) with the current estimates. The exact form of the CDA updates are given in the subsequent section. Upon convergence of IRLS, we update $\hat{\phi}_j$ once via Newton-Raphson (Piegorsch, 1990). Specifically, we estimate $\hat{\phi}_j$ via the `MASS` R package (Venables and Ripley, 2002).

Convergence of the IRLS is determined by a threshold on the mean absolute relative change of parameter estimates across one IRLS iteration. In particular, let $(r)$ index the current IRLS iteration, such that $\hat{\boldsymbol{\Theta}}_j^{(r)} = (\hat{\boldsymbol{\beta}}_j^{(r)}, \hat{\boldsymbol{\gamma}}_j^{(r)}) = (\hat{\beta}_{j1}^{(r)}, \ldots, \hat{\beta}_{jK}^{(r)}, \hat{\gamma}_{j1}^{(r)}, \ldots, \hat{\gamma}_{jP}^{(r)})$ denote the estimates of the parameters $\boldsymbol{\beta}_j$ and $\boldsymbol{\gamma}_j$ at the end of the current $r^{th}$ IRLS iteration. Then, for $\Theta_{jt}$ denoting the $t^{th}$ element of $\boldsymbol{\Theta}_j$, convergence of the IRLS is attained when:

$$\frac{1}{K+P}\sum_{t=1}^{K+P}\left|\frac{\hat{\Theta}_{jt}^{(r)} - \hat{\Theta}_{jt}^{(r-1)}}{\hat{\Theta}_{jt}^{(r-1)}}\right| < \epsilon_2.$$

In FSCseq, we set $\epsilon_2 = 10^{-4}$ as the default convergence threshold. The details and convergence criterion for the embedded coordinate-descent algorithm (CDA) can be found in Section A3.

#### A3: Coordinate-wise Descent Updates

The coordinate-wise descent algorithm (CDA) updates for $(\theta_{j,kl}, \beta_{jk}, \gamma_{jp})$ for each gene $j = 1, \ldots, G$, clusters $1 \leq k < l \leq K$, and covariates $p = 1, \ldots, P$ are given in this section. Current estimates of parameters are denoted by tilde, e.g. $\tilde{\beta}_{jk}$, and the new CDA update is denoted by a hat, e.g. $\hat{\beta}_{jk}$. Let $(m+1)$ be the current iteration index of the M-step in the EM/CEM algorithm. Then, in the first IRLS iteration, the values of the current estimates from the previous M step update are initialized in CDA as $(\tilde{\beta}_{jk}, \tilde{\gamma}_{jp}) \leftarrow (\hat{\beta}_{jk}^{(m)}, \hat{\gamma}_{jp}^{(m)})$. Subsequent CDA loops are initialized with the most recent CDA updates. For clarity of notation, let $v_{ij}$ denote the $i^{th}$ element of the diagonal entries of $\mathbf{W}_j$ for all $i = 1, \ldots, N$ with $N = n \cdot K$. Then, the CDA update equations for fixed gene $j$ are given as follows:

$$\hat{\theta}_{j,kl} = \begin{cases} sgn(\tilde{\theta}_{j,kl})\left(\left|\tilde{\theta}_{j,kl}\right| - \frac{\alpha}{1-\alpha}\right)_+, & \left|\tilde{\theta}_{j,kl}\right| \leq \lambda^* \\ sgn(\tilde{\omega}_{j,kl})\left(\left|\tilde{\omega}_{j,kl}\right| - \frac{a\frac{\alpha}{1-\alpha}}{a-1-\frac{1}{\lambda(1-\alpha)}}\right)_+, & \lambda^* < \left|\tilde{\theta}_{j,kl}\right| \leq a\lambda\alpha \\ \tilde{\theta}_{j,kl}, & \left|\tilde{\theta}_{j,kl}\right| > a\lambda\alpha \end{cases} \tag{3}$$

$$\hat{\beta}_{jk} = \frac{\frac{1}{n_k}\sum_{i=1}^N v_{ij}x_{ik}\tilde{r}_{ijk} + \lambda(1-\alpha)[\sum_{l>k}(\tilde{\beta}_{jl} + \tilde{\theta}_{j,kl}) + \sum_{l<k}(\tilde{\beta}_{jl} - \tilde{\theta}_{j,lk})]}{\lambda(1-\alpha)(K-1) + \frac{1}{n_k}\sum_{i=1}^N v_{ij}x_{ik}^2}$$

$$\hat{\gamma}_{jp} = \frac{\sum_{i=1}^N v_{ij}x_{ip}\tilde{r}_{ijp}}{\sum_{i=1}^N v_{ij}x_{ip}^2}$$

where $(a)_+$ is equal to $a$ if $a > 0$ and is equal to 0 otherwise, and $x_{ik}$ is the $i^{th}$ row and $k^{th}$ column entry of the design matrix $\mathbf{X}$ as defined in Section A2. The effective number of observations used for estimation of parameters is $N = n \cdot K$, denoting each of the IRLS-weighted $n$ samples in each of the $K$ clusters. The estimated number of samples in cluster $k$ (at the current M-step) used for estimation of $\beta_{jk}$ is $n_k = \hat{\pi}_k^{(m+1)} n$. Also, $\tilde{r}_{ijk} = \tilde{y}_{ijk} - \sum_{c \neq k}^{K} x_{ic}\tilde{\beta}_{jc} - \sum_{p=1}^{P} x_{ip}\tilde{\gamma}_{jp}$; $\tilde{r}_{ijp} = \tilde{y}_{ijk} - \sum_{c=1}^{K} x_{ic}\tilde{\beta}_{jc} - \sum_{q \neq p}^{P} x_{iq}\tilde{\gamma}_{jq}$; $\tilde{\omega}_{j,kl} = \frac{(a-1)\tilde{\theta}_{j,kl}}{a-1-\frac{1}{\lambda(1-\alpha)}}$; and $\lambda^* = \dfrac{\alpha}{1-\alpha} + \lambda\alpha$.

We iteratively update these parameters until convergence of CDA (inner loop) for fixed gene $j$. Then, we recompute the IRLS weights and repeat, until the IRLS (outer loop) converges. The dispersion estimate $\hat{\phi}_j$ is attained upon convergence of IRLS, as described in Section A2. Then, we output all of the final updates as the estimates from the current $(m+1)^{th}$ M step, i.e. $(\hat{\beta}_{jk}^{(m+1)}, \hat{\gamma}_{jp}^{(m+1)}, \hat{\pi}_k^{(m+1)}, \hat{\phi}_j^{(m+1)}) \leftarrow (\hat{\beta}_{jk}, \hat{\gamma}_{jp}, \hat{\pi}_k, \hat{\phi}_j)$. This entire procedure is repeated for each gene $j = 1, \ldots, G$, or if mini-batching, $j \in minibatch^{(m+1)}$, where $minibatch^{(m+1)}$ is the mini-batch of genes that were selected at the current M step. We note that estimation of $\hat{\pi}_k$ is done before the IRLS, as described in Algorithm 1 of the main text.

Convergence of the CDA (within each IRLS iteration) is attained when the mean absolute relative change of the parameter estimates becomes small, similar to the IRLS. In particular, let $\hat{\mathbf{\Theta}}_j = (\hat{\boldsymbol{\beta}}_j, \hat{\boldsymbol{\gamma}}_j) = (\hat{\beta}_{j1}, \ldots, \hat{\beta}_{jK}, \hat{\gamma}_{j1}, \ldots, \hat{\gamma}_{jP})$ denote the estimates of the parameters $\boldsymbol{\beta}_j$ and $\boldsymbol{\gamma}_j$ at the end of the current CDA iteration, and let $\tilde{\mathbf{\Theta}}_j$ similarly denote the corresponding estimates from the end of the previous CDA iteration. Then, for $\Theta_{jt}$ denoting the $t^{th}$ element of $\mathbf{\Theta}_j$, convergence of the CDA is attained when

$$\frac{1}{K+P} \sum_{t=1}^{K+P} \left| \frac{\hat{\Theta}_{jt} - \tilde{\Theta}_{jt}}{\tilde{\Theta}_{jt}} \right| < \epsilon_2.$$

By default, we set $\epsilon_2$ to be the same for the convergence criteria of the CDA and the IRLS. Importantly, for each CDA iteration, convergence is checked once at the end of the iteration (i.e., when all parameters $\mathbf{\Theta}_j$ have been updated within that iteration)

### Section B: Performance of FSCseq on all simulated conditions

In this section, we show all results from running the default FSCseq scheme on our extensive set of simulation cases. Supplementary Tables 1 and 2 show all results from every combination of simulated conditions, averaged over 25 simulation runs each.

| **n** | **LFC** | $\mathbf{p_{DE}}$ | $\boldsymbol{\beta_0}$ | $\boldsymbol{\phi_0}$ | $\bar{K}^*$ | $OA$ | $\overline{ARI}$ | $\overline{pARI}$ | $\overline{TPR}$ | $\overline{FPR}$ |
|---|---|---|---|---|---|---|---|---|---|---|
| 50 | 1.0 | 0.025 | 8 | 0.15 | 2.00 | 1.00 | 1.00 | 1.00 | 0.946 | 0.0005 |
| | | | | 0.35 | 2.00 | 1.00 | 1.00 | 1.00 | 0.635 | 0.0004 |
| | | | | 0.50 | 2.00 | 1.00 | 1.00 | 1.00 | 0.413 | 0.0004 |
| | | | 12 | 0.15 | 2.00 | 1.00 | 1.00 | 1.00 | 0.952 | 0.0003 |
| | | | | 0.35 | 2.00 | 1.00 | 1.00 | 1.00 | 0.624 | 0.0004 |
| | | | | 0.50 | 2.00 | 1.00 | 1.00 | 1.00 | 0.397 | 0.0004 |
| | | 0.050 | 8 | 0.15 | 2.00 | 1.00 | 1.00 | 1.00 | 0.958 | 0.0005 |
| | | | | 0.35 | 2.08 | 0.92 | 0.99 | 1.00 | 0.654 | 0.0005 |
| | | | | 0.50 | 2.00 | 1.00 | 1.00 | 1.00 | 0.414 | 0.0004 |
| | | | 12 | 0.15 | 2.00 | 1.00 | 1.00 | 1.00 | 0.956 | 0.0003 |
| | | | | 0.35 | 2.04 | 0.96 | 0.99 | 1.00 | 0.666 | 0.0003 |
| | | | | 0.50 | 2.00 | 1.00 | 1.00 | 1.00 | 0.418 | 0.0004 |
| | 2.0 | 0.025 | 8 | 0.15 | 2.00 | 1.00 | 1.00 | 1.00 | 0.983 | 0.0005 |
| | | | | 0.35 | 2.00 | 1.00 | 1.00 | 1.00 | 0.958 | 0.0004 |
| | | | | 0.50 | 2.00 | 1.00 | 1.00 | 1.00 | 0.958 | 0.0004 |
| | | | 12 | 0.15 | 2.00 | 1.00 | 1.00 | 1.00 | 0.992 | 0.0003 |
| | | | | 0.35 | 2.04 | 0.96 | 0.99 | 0.99 | 0.967 | 0.0003 |
| | | | | 0.50 | 2.00 | 1.00 | 1.00 | 1.00 | 0.957 | 0.0005 |
| | | 0.050 | 8 | 0.15 | 2.00 | 1.00 | 1.00 | 1.00 | 0.998 | 0.0004 |
| | | | | 0.35 | 2.00 | 1.00 | 1.00 | 1.00 | 0.973 | 0.0005 |
| | | | | 0.50 | 2.04 | 0.96 | 1.00 | 1.00 | 0.955 | 0.0004 |
| | | | 12 | 0.15 | 2.04 | 0.96 | 1.00 | 1.00 | 0.999 | 0.0002 |
| | | | | 0.35 | 2.00 | 1.00 | 1.00 | 1.00 | 0.982 | 0.0004 |
| | | | | 0.50 | 2.00 | 1.00 | 1.00 | 1.00 | 0.960 | 0.0004 |
| 100 | 1.0 | 0.025 | 8 | 0.15 | 2.00 | 1.00 | 1.00 | 1.00 | 0.952 | 0.0003 |
| | | | | 0.35 | 2.00 | 1.00 | 1.00 | 1.00 | 0.924 | 0.0004 |
| | | | | 0.50 | 2.00 | 1.00 | 1.00 | 1.00 | 0.835 | 0.0003 |
| | | | 12 | 0.15 | 2.00 | 1.00 | 1.00 | 1.00 | 0.959 | 0.0002 |
| | | | | 0.35 | 2.00 | 1.00 | 1.00 | 1.00 | 0.919 | 0.0004 |
| | | | | 0.50 | 2.00 | 1.00 | 1.00 | 1.00 | 0.836 | 0.0003 |
| | | 0.050 | 8 | 0.15 | 2.00 | 1.00 | 1.00 | 1.00 | 0.968 | 0.0002 |
| | | | | 0.35 | 2.00 | 1.00 | 1.00 | 1.00 | 0.921 | 0.0004 |
| | | | | 0.50 | 2.00 | 1.00 | 1.00 | 1.00 | 0.833 | 0.0003 |
| | | | 12 | 0.15 | 2.00 | 1.00 | 1.00 | 1.00 | 0.974 | 0.0003 |
| | | | | 0.35 | 2.00 | 1.00 | 1.00 | 1.00 | 0.923 | 0.0005 |
| | | | | 0.50 | 2.00 | 1.00 | 1.00 | 1.00 | 0.838 | 0.0003 |
| | 2.0 | 0.025 | 8 | 0.15 | 2.00 | 1.00 | 1.00 | 1.00 | 0.989 | 0.0002 |
| | | | | 0.35 | 2.00 | 1.00 | 1.00 | 1.00 | 0.963 | 0.0003 |
| | | | | 0.50 | 2.00 | 1.00 | 1.00 | 1.00 | 0.950 | 0.0003 |
| | | | 12 | 0.15 | 2.00 | 1.00 | 1.00 | 1.00 | 0.988 | 0.0003 |
| | | | | 0.35 | 2.00 | 1.00 | 1.00 | 1.00 | 0.971 | 0.0003 |
| | | | | 0.50 | 2.00 | 1.00 | 1.00 | 1.00 | 0.963 | 0.0003 |
| | | 0.050 | 8 | 0.15 | 2.00 | 1.00 | 1.00 | 1.00 | 0.997 | 0.0002 |
| | | | | 0.35 | 2.00 | 1.00 | 1.00 | 1.00 | 0.975 | 0.0003 |
| | | | | 0.50 | 2.00 | 1.00 | 1.00 | 1.00 | 0.969 | 0.0002 |
| | | | 12 | 0.15 | 2.00 | 1.00 | 1.00 | 1.00 | 0.998 | 0.0002 |
| | | | | 0.35 | 2.00 | 1.00 | 1.00 | 1.00 | 0.981 | 0.0002 |
| | | | | 0.50 | 2.00 | 1.00 | 1.00 | 1.00 | 0.966 | 0.0003 |

Supplementary Table 1: FSCseq results from all simulation conditions with $K_{true} = 2$ underlying groups.

| n | LFC | $p_{DE}$ | $\beta_0$ | $\phi_0$ | $\bar{K}^*$ | $OA$ | $\overline{ARI}$ | $\overline{pARI}$ | $\overline{TPR}$ | $\overline{FPR}$ |
|---|---|---|---|---|---|---|---|---|---|---|
| 100 | 1.0 | 0.025 | 8 | 0.15 | 4.00 | 0.92 | 0.99 | 0.99 | 0.695 | 0.0004 |
| | | | | 0.35 | 3.96 | 0.80 | 0.96 | 0.95 | 0.583 | 0.0014 |
| | | | | 0.50 | 2.20 | 0.00 | 0.24 | 0.21 | 0.079 | 0.0002 |
| | | | 12 | 0.15 | 4.08 | 0.84 | 0.98 | 0.99 | 0.683 | 0.0011 |
| | | | | 0.35 | 3.96 | 0.68 | 0.93 | 0.92 | 0.540 | 0.0015 |
| | | | | 0.50 | 2.32 | 0.04 | 0.24 | 0.20 | 0.078 | 0.0005 |
| | | 0.050 | 8 | 0.15 | 4.08 | 0.92 | 0.99 | 1.00 | 0.671 | 0.0012 |
| | | | | 0.35 | 3.96 | 0.96 | 0.99 | 0.99 | 0.610 | 0.0016 |
| | | | | 0.50 | 3.92 | 0.92 | 0.98 | 0.97 | 0.448 | 0.0014 |
| | | | 12 | 0.15 | 4.00 | 1.00 | 1.00 | 1.00 | 0.687 | 0.0004 |
| | | | | 0.35 | 3.96 | 0.96 | 0.99 | 0.99 | 0.611 | 0.0018 |
| | | | | 0.50 | 3.76 | 0.80 | 0.94 | 0.92 | 0.448 | 0.0011 |
| | 2.0 | 0.025 | 8 | 0.15 | 4.12 | 0.88 | 0.99 | 1.00 | 0.762 | 0.0012 |
| | | | | 0.35 | 4.00 | 1.00 | 1.00 | 1.00 | 0.719 | 0.0001 |
| | | | | 0.50 | 4.00 | 1.00 | 1.00 | 1.00 | 0.726 | 0.0014 |
| | | | 12 | 0.15 | 4.04 | 0.96 | 1.00 | 1.00 | 0.773 | 0.0002 |
| | | | | 0.35 | 4.00 | 1.00 | 1.00 | 1.00 | 0.748 | 0.0002 |
| | | | | 0.50 | 4.00 | 1.00 | 1.00 | 1.00 | 0.724 | 0.0011 |
| | | 0.050 | 8 | 0.15 | 4.12 | 0.88 | 1.00 | 1.00 | 0.756 | 0.0017 |
| | | | | 0.35 | 4.04 | 0.96 | 1.00 | 1.00 | 0.743 | 0.0008 |
| | | | | 0.50 | 4.00 | 1.00 | 1.00 | 1.00 | 0.717 | 0.0012 |
| | | | 12 | 0.15 | 4.00 | 1.00 | 1.00 | 1.00 | 0.775 | 0.0000 |
| | | | | 0.35 | 4.08 | 0.92 | 0.99 | 1.00 | 0.750 | 0.0020 |
| | | | | 0.50 | 4.04 | 0.96 | 1.00 | 1.00 | 0.727 | 0.0025 |
| 200 | 1.0 | 0.025 | 8 | 0.15 | 4.28 | 0.72 | 0.98 | 1.00 | 0.722 | 0.0017 |
| | | | | 0.35 | 4.12 | 0.84 | 0.98 | 1.00 | 0.665 | 0.0007 |
| | | | | 0.50 | 4.08 | 0.92 | 0.99 | 1.00 | 0.640 | 0.0010 |
| | | | 12 | 0.15 | 4.16 | 0.84 | 0.99 | 1.00 | 0.744 | 0.0006 |
| | | | | 0.35 | 3.96 | 0.96 | 0.99 | 0.99 | 0.678 | 0.0009 |
| | | | | 0.50 | 4.00 | 0.92 | 0.98 | 0.98 | 0.670 | 0.0018 |
| | | 0.050 | 8 | 0.15 | 4.12 | 0.88 | 1.00 | 1.00 | 0.718 | 0.0009 |
| | | | | 0.35 | 4.00 | 1.00 | 1.00 | 1.00 | 0.689 | 0.0028 |
| | | | | 0.50 | 4.12 | 0.88 | 1.00 | 1.00 | 0.673 | 0.0027 |
| | | | 12 | 0.15 | 4.28 | 0.72 | 0.98 | 0.99 | 0.720 | 0.0030 |
| | | | | 0.35 | 4.00 | 1.00 | 1.00 | 1.00 | 0.709 | 0.0038 |
| | | | | 0.50 | 4.00 | 1.00 | 1.00 | 1.00 | 0.666 | 0.0013 |
| | 2.0 | 0.025 | 8 | 0.15 | 4.08 | 0.92 | 0.99 | 1.00 | 0.812 | 0.0012 |
| | | | | 0.35 | 4.00 | 1.00 | 1.00 | 1.00 | 0.803 | 0.0001 |
| | | | | 0.50 | 4.16 | 0.84 | 0.99 | 1.00 | 0.763 | 0.0011 |
| | | | 12 | 0.15 | 4.08 | 0.92 | 0.99 | 1.00 | 0.797 | 0.0017 |
| | | | | 0.35 | 4.16 | 0.84 | 1.00 | 1.00 | 0.781 | 0.0009 |
| | | | | 0.50 | 4.16 | 0.84 | 0.99 | 1.00 | 0.781 | 0.0004 |
| | | 0.050 | 8 | 0.15 | 4.04 | 0.96 | 1.00 | 1.00 | 0.822 | 0.0003 |
| | | | | 0.35 | 4.28 | 0.72 | 0.99 | 0.99 | 0.777 | 0.0063 |
| | | | | 0.50 | 4.00 | 1.00 | 1.00 | 1.00 | 0.772 | 0.0001 |
| | | | 12 | 0.15 | 4.04 | 0.96 | 1.00 | 1.00 | 0.806 | 0.0005 |
| | | | | 0.35 | 4.00 | 1.00 | 1.00 | 1.00 | 0.767 | 0.0000 |
| | | | | 0.50 | 4.04 | 0.96 | 1.00 | 1.00 | 0.776 | 0.0002 |

Supplementary Table 2: FSCseq results from all simulation conditions with $K_{true} = 4$ underlying groups.

Additionally, we show Figure 1 of the main text without filtering outlier points.

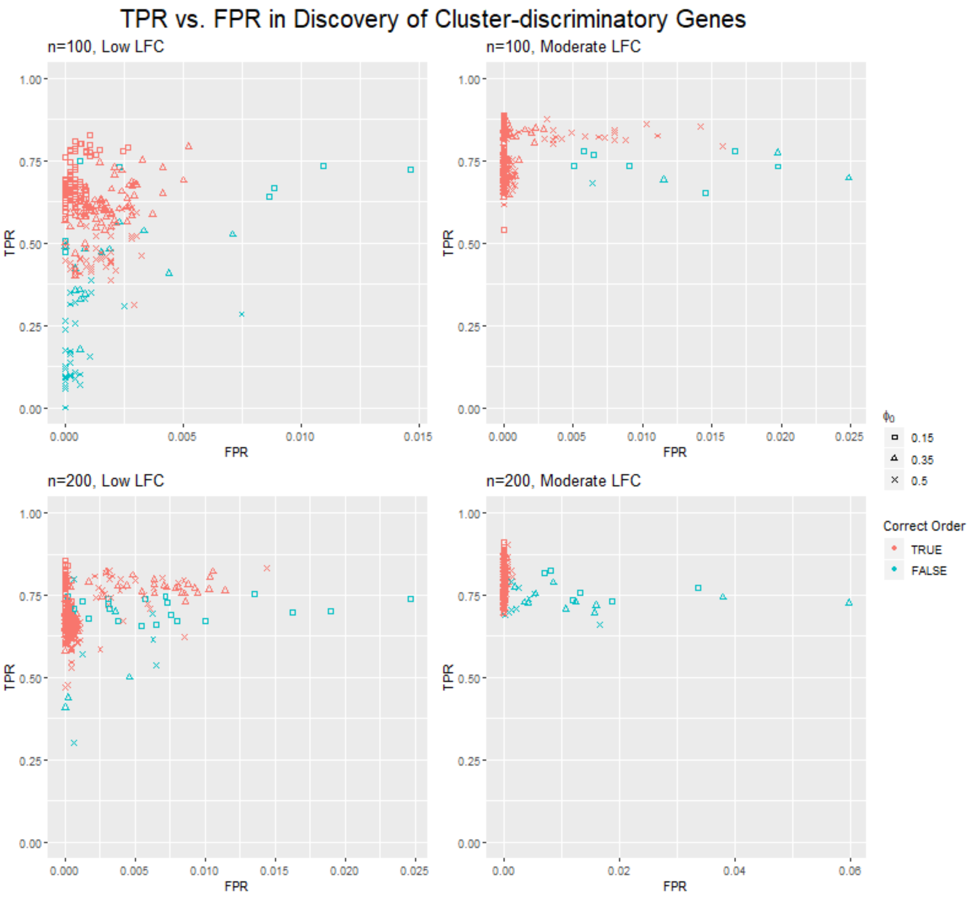


Supplementary Figure 2: Plots corresponding to Figure 1 of the main text, without removing outlier points. Scatterplot of true positive rate ($TPR$) vs false positive rate ($FPR$) in discovering cluster-discriminatory genes in simulated datasets via FSCseq. Displayed points correspond to simulated datasets with $K_{true} = 4$ underlying clusters, with $n = 100$ (top) or 200 (bottom) and simulated $LFC = 1$ ('Low', left) or 2 ('Moderate', right). Red points indicate that the correct order was uncovered ($K^* = K_{true}$), and blue points indicate that an incorrect order was uncovered. Squares, circles, and X's indicate low ($\phi_0 = 0.15$), moderate ($\phi_0 = 0.35$), and high ($\phi_0 = 0.50$) levels of overdispersion, respectively.

### Section C: Comparative Methods

#### C1: Automation of iCluster+ order selection

In order to determine the optimal number of clusters $K^*$, the `iCluster+` paper and manual suggest searching graphically for the value of $K$ at which the deviance ratio (% Variability Explained) hits a plateau. This is necessary because the deviance ratio will continue to increase with higher order, for increased noise in the dataset (Mo et al., 2013; Mo and Shen, 2019). We observed this pattern even at very low levels of overdispersion, causing `iCluster+`

to consistently attain the maximum deviance ratio at the largest candidate value of $K$. In searching extensive sets of simulations, it becomes infeasible to manually/visually search for optimal parameters. Thus, to systemize the procedure across many simulation cases, we selected an arbitrary threshold based on the manual, such that if the percent increase in variability explained is less than 0.05 with an added cluster, the optimal $K^*$ is selected as the immediately previous value of $K$.

#### C2: Clustering performance on simulated datasets

In this section, we show the performance of FSCseq and competing methods across all simulated conditions. Supplementary Tables 3 and 4, and Supplementary Tables 5 and 6 show the average discovered order ($\bar{K}^*$) and average attained ARI ($\overline{ARI}$) for all simulation conditions with $K_{true} = 2$ and $K_{true} = 4$, respectively. For each row, the best performing method's metric is colored in red.

| n | LFC | $p_{DE}$ | $\beta_0$ | $\phi_0$ | $\bar{K}^*_{FSC}$ | $\bar{K}^*_{iCl}$ | $\bar{K}^*_{HC}$ | $\bar{K}^*_{KM}$ | $\bar{K}^*_{NBMB}$ | $\bar{K}^*_{lMC}$ | $\bar{K}^*_{vMC}$ | $\bar{K}^*_{rMC}$ |
|---|---|---|---|---|---|---|---|---|---|---|---|---|
| 50 | 1.0 | 0.025 | 8 | 0.15 | 2.00 | 6.00 | 2.00 | 2.00 | 2.00 | 2.00 | 2.00 | 2.00 |
| | | | | 0.35 | 2.00 | 6.00 | 2.00 | 2.00 | 2.00 | 2.00 | 2.00 | 2.00 |
| | | | | 0.50 | 2.00 | 6.00 | 2.00 | 2.32 | 2.00 | 2.00 | 2.00 | 2.00 |
| | | | 12 | 0.15 | 2.00 | 6.00 | 2.00 | 2.00 | 2.00 | 2.00 | 2.00 | 2.00 |
| | | | | 0.35 | 2.00 | 6.00 | 2.00 | 2.04 | 2.00 | 2.00 | 2.00 | 2.00 |
| | | | | 0.50 | 2.00 | 6.00 | 2.00 | 2.20 | 2.00 | 2.00 | 2.00 | 2.00 |
| | | 0.050 | 8 | 0.15 | 2.00 | 6.00 | 2.00 | 2.00 | 2.36 | 2.00 | 2.00 | 2.00 |
| | | | | 0.35 | 2.08 | 6.00 | 2.00 | 2.00 | 2.00 | 2.00 | 2.00 | 2.00 |
| | | | | 0.50 | 2.00 | 6.00 | 2.00 | 2.04 | 2.00 | 2.00 | 2.00 | 2.00 |
| | | | 12 | 0.15 | 2.00 | 6.00 | 2.00 | 2.00 | 2.72 | 2.00 | 2.00 | 2.00 |
| | | | | 0.35 | 2.04 | 6.00 | 2.00 | 2.00 | 2.00 | 2.00 | 2.00 | 2.00 |
| | | | | 0.50 | 2.00 | 6.00 | 2.00 | 2.00 | 2.00 | 2.00 | 2.00 | 2.00 |
| | 2.0 | 0.025 | 8 | 0.15 | 2.00 | 6.00 | 2.00 | 2.00 | 2.64 | 2.00 | 2.00 | 2.00 |
| | | | | 0.35 | 2.00 | 6.00 | 2.00 | 2.00 | 2.00 | 2.00 | 2.00 | 2.00 |
| | | | | 0.50 | 2.00 | 6.00 | 2.00 | 2.00 | 2.00 | 2.00 | 2.00 | 2.00 |
| | | | 12 | 0.15 | 2.00 | 6.00 | 2.00 | 2.00 | 2.36 | 2.00 | 2.00 | 2.00 |
| | | | | 0.35 | 2.04 | 6.00 | 2.00 | 2.00 | 2.24 | 2.00 | 2.00 | 2.00 |
| | | | | 0.50 | 2.00 | 6.00 | 2.00 | 2.00 | 2.04 | 2.00 | 2.00 | 2.00 |
| | | 0.050 | 8 | 0.15 | 2.00 | 6.00 | 2.00 | 2.00 | 2.36 | 2.00 | 2.00 | 2.00 |
| | | | | 0.35 | 2.00 | 6.00 | 2.00 | 2.00 | 2.16 | 2.00 | 2.00 | 2.00 |
| | | | | 0.50 | 2.04 | 6.00 | 2.00 | 2.00 | 2.24 | 2.00 | 2.00 | 2.00 |
| | | | 12 | 0.15 | 2.04 | 6.00 | 2.00 | 2.00 | 2.64 | 2.00 | 2.00 | 2.00 |
| | | | | 0.35 | 2.00 | 6.00 | 2.00 | 2.00 | 2.56 | 2.00 | 2.00 | 2.00 |
| | | | | 0.50 | 2.00 | 6.00 | 2.00 | 2.00 | 2.64 | 2.00 | 2.00 | 2.00 |
| 100 | 1.0 | 0.025 | 8 | 0.15 | 2.00 | 6.00 | 2.00 | 2.00 | 3.00 | 2.00 | 2.00 | 2.00 |
| | | | | 0.35 | 2.00 | 6.00 | 2.00 | 2.00 | 2.00 | 2.00 | 2.00 | 2.00 |
| | | | | 0.50 | 2.00 | 6.00 | 2.00 | 2.04 | 2.00 | 2.00 | 2.00 | 2.00 |
| | | | 12 | 0.15 | 2.00 | 6.00 | 2.00 | 2.00 | 2.96 | 2.00 | 2.00 | 2.00 |
| | | | | 0.35 | 2.00 | 6.00 | 2.00 | 2.00 | 2.00 | 2.00 | 2.00 | 2.00 |
| | | | | 0.50 | 2.00 | 6.00 | 2.00 | 2.04 | 2.00 | 2.00 | 2.00 | 2.00 |
| | | 0.050 | 8 | 0.15 | 2.00 | 6.00 | 2.00 | 2.00 | 2.80 | 2.00 | 2.00 | 2.00 |
| | | | | 0.35 | 2.00 | 6.00 | 2.00 | 2.00 | 3.28 | 2.00 | 2.00 | 2.00 |
| | | | | 0.50 | 2.00 | 6.00 | 2.00 | 2.00 | 2.28 | 2.00 | 2.00 | 2.00 |
| | | | 12 | 0.15 | 2.00 | 6.00 | 2.00 | 2.00 | 3.24 | 2.00 | 2.00 | 2.00 |
| | | | | 0.35 | 2.00 | 6.00 | 2.00 | 2.00 | 3.32 | 2.00 | 2.00 | 2.00 |
| | | | | 0.50 | 2.00 | 6.00 | 2.00 | 2.04 | 2.20 | 2.00 | 2.00 | 2.00 |
| | 2.0 | 0.025 | 8 | 0.15 | 2.00 | 6.00 | 2.00 | 2.00 | 2.48 | 2.00 | 2.00 | 2.00 |
| | | | | 0.35 | 2.00 | 6.00 | 2.00 | 2.00 | 2.80 | 2.00 | 2.00 | 2.00 |
| | | | | 0.50 | 2.00 | 6.00 | 2.00 | 2.00 | 2.52 | 2.00 | 2.00 | 2.00 |
| | | | 12 | 0.15 | 2.00 | 6.00 | 2.00 | 2.00 | 3.40 | 2.00 | 2.00 | 2.00 |
| | | | | 0.35 | 2.00 | 6.00 | 2.00 | 2.00 | 4.08 | 2.00 | 2.00 | 2.00 |
| | | | | 0.50 | 2.00 | 6.00 | 2.00 | 2.00 | 4.16 | 2.00 | 2.00 | 2.00 |
| | | 0.050 | 8 | 0.15 | 2.00 | 2.00 | 2.00 | 2.00 | 2.40 | 2.00 | 2.00 | 2.00 |
| | | | | 0.35 | 2.00 | 6.00 | 2.00 | 2.00 | 2.24 | 2.00 | 2.00 | 2.00 |
| | | | | 0.50 | 2.00 | 6.00 | 2.00 | 2.00 | 2.72 | 2.00 | 2.00 | 2.00 |
| | | | 12 | 0.15 | 2.00 | 2.00 | 2.00 | 2.00 | 2.16 | 2.00 | 2.00 | 2.00 |
| | | | | 0.35 | 2.00 | 6.00 | 2.00 | 2.00 | 3.12 | 2.00 | 2.00 | 2.00 |
| | | | | 0.50 | 2.00 | 6.00 | 2.00 | 2.00 | 3.92 | 2.00 | 2.00 | 2.00 |

Supplementary Table 3: Average estimated orders $\bar{K}^*$ from compared methods for all simulation conditions (rows) with $K_{true} = 2$ underlying groups. Best performance within each row is colored in red.

| n | LFC | $p_{DE}$ | $\beta_0$ | $\phi_0$ | $\overline{ARI}_{FSC}$ | $\overline{ARI}_{iCl}$ | $\overline{ARI}_{HC}$ | $\overline{ARI}_{KM}$ | $\overline{ARI}_{NBMB}$ | $\overline{ARI}_{lMC}$ | $\overline{ARI}_{vMC}$ | $\overline{ARI}_{rMC}$ |
|---|---|---|---|---|---|---|---|---|---|---|---|---|
| 50 | 1.0 | 0.025 | 8 | 0.15 | 1.00 | 0.27 | 1.00 | 1.00 | 0.17 | 1.00 | 1.00 | 1.00 |
| | | | | 0.35 | 1.00 | 0.26 | 0.89 | 0.50 | 0.04 | 0.87 | 0.89 | 0.88 |
| | | | | 0.50 | 1.00 | 0.24 | 0.74 | 0.23 | 0.03 | 0.67 | 0.78 | 0.71 |
| | | | 12 | 0.15 | 1.00 | 0.32 | 1.00 | 0.99 | 0.10 | 1.00 | 1.00 | 1.00 |
| | | | | 0.35 | 1.00 | 0.28 | 0.84 | 0.53 | 0.01 | 0.89 | 0.86 | 0.87 |
| | | | | 0.50 | 1.00 | 0.26 | 0.78 | 0.24 | 0.03 | 0.68 | 0.70 | 0.70 |
| | | 0.050 | 8 | 0.15 | 1.00 | 0.32 | 1.00 | 1.00 | 0.34 | 1.00 | 1.00 | 1.00 |
| | | | | 0.35 | 0.99 | 0.30 | 0.99 | 0.93 | 0.14 | 1.00 | 1.00 | 1.00 |
| | | | | 0.50 | 1.00 | 0.28 | 0.97 | 0.61 | 0.05 | 0.92 | 0.96 | 0.95 |
| | | | 12 | 0.15 | 1.00 | 0.32 | 1.00 | 1.00 | 0.25 | 1.00 | 1.00 | 1.00 |
| | | | | 0.35 | 0.99 | 0.31 | 1.00 | 0.92 | 0.16 | 1.00 | 1.00 | 1.00 |
| | | | | 0.50 | 1.00 | 0.28 | 0.97 | 0.62 | 0.09 | 0.94 | 0.95 | 0.95 |
| | 2.0 | 0.025 | 8 | 0.15 | 1.00 | 0.32 | 1.00 | 1.00 | 0.63 | 1.00 | 1.00 | 1.00 |
| | | | | 0.35 | 1.00 | 0.30 | 1.00 | 1.00 | 0.48 | 1.00 | 1.00 | 0.99 |
| | | | | 0.50 | 1.00 | 0.30 | 1.00 | 0.88 | 0.14 | 0.97 | 0.98 | 0.97 |
| | | | 12 | 0.15 | 1.00 | 0.34 | 1.00 | 1.00 | 0.16 | 1.00 | 1.00 | 1.00 |
| | | | | 0.35 | 0.99 | 0.32 | 0.99 | 1.00 | 0.17 | 1.00 | 1.00 | 1.00 |
| | | | | 0.50 | 1.00 | 0.32 | 1.00 | 1.00 | 0.16 | 1.00 | 1.00 | 1.00 |
| | | 0.050 | 8 | 0.15 | 1.00 | 0.35 | 1.00 | 1.00 | 0.88 | 1.00 | 1.00 | 1.00 |
| | | | | 0.35 | 1.00 | 0.33 | 1.00 | 1.00 | 0.92 | 1.00 | 1.00 | 1.00 |
| | | | | 0.50 | 1.00 | 0.32 | 1.00 | 1.00 | 0.71 | 1.00 | 1.00 | 1.00 |
| | | | 12 | 0.15 | 1.00 | 0.35 | 1.00 | 1.00 | 0.69 | 1.00 | 1.00 | 1.00 |
| | | | | 0.35 | 1.00 | 0.34 | 1.00 | 1.00 | 0.49 | 1.00 | 1.00 | 1.00 |
| | | | | 0.50 | 1.00 | 0.33 | 1.00 | 1.00 | 0.45 | 1.00 | 1.00 | 1.00 |
| 100 | 1.0 | 0.025 | 8 | 0.15 | 1.00 | 0.30 | 1.00 | 1.00 | 0.29 | 1.00 | 1.00 | 1.00 |
| | | | | 0.35 | 1.00 | 0.30 | 0.91 | 0.55 | 0.22 | 0.97 | 0.95 | 0.95 |
| | | | | 0.50 | 1.00 | 0.29 | 0.76 | 0.23 | 0.10 | 0.73 | 0.75 | 0.78 |
| | | | 12 | 0.15 | 1.00 | 0.34 | 1.00 | 1.00 | 0.40 | 1.00 | 1.00 | 1.00 |
| | | | | 0.35 | 1.00 | 0.33 | 0.93 | 0.59 | 0.11 | 0.97 | 0.97 | 0.93 |
| | | | | 0.50 | 1.00 | 0.30 | 0.78 | 0.22 | 0.14 | 0.73 | 0.75 | 0.79 |
| | | 0.050 | 8 | 0.15 | 1.00 | 0.33 | 1.00 | 1.00 | 0.76 | 1.00 | 1.00 | 1.00 |
| | | | | 0.35 | 1.00 | 0.33 | 1.00 | 0.96 | 0.41 | 1.00 | 1.00 | 1.00 |
| | | | | 0.50 | 1.00 | 0.33 | 0.98 | 0.69 | 0.35 | 0.99 | 0.99 | 1.00 |
| | | | 12 | 0.15 | 1.00 | 0.35 | 1.00 | 1.00 | 0.69 | 1.00 | 1.00 | 1.00 |
| | | | | 0.35 | 1.00 | 0.34 | 1.00 | 0.95 | 0.44 | 1.00 | 1.00 | 1.00 |
| | | | | 0.50 | 1.00 | 0.33 | 0.98 | 0.68 | 0.36 | 1.00 | 1.00 | 1.00 |
| | 2.0 | 0.025 | 8 | 0.15 | 1.00 | 0.33 | 1.00 | 1.00 | 0.84 | 1.00 | 1.00 | 1.00 |
| | | | | 0.35 | 1.00 | 0.33 | 1.00 | 1.00 | 0.57 | 1.00 | 1.00 | 1.00 |
| | | | | 0.50 | 1.00 | 0.33 | 1.00 | 0.91 | 0.27 | 1.00 | 1.00 | 0.99 |
| | | | 12 | 0.15 | 1.00 | 0.34 | 1.00 | 1.00 | 0.60 | 1.00 | 1.00 | 1.00 |
| | | | | 0.35 | 1.00 | 0.34 | 1.00 | 1.00 | 0.38 | 1.00 | 1.00 | 1.00 |
| | | | | 0.50 | 1.00 | 0.34 | 1.00 | 0.99 | 0.21 | 1.00 | 1.00 | 1.00 |
| | | 0.050 | 8 | 0.15 | 1.00 | 1.00 | 1.00 | 1.00 | 0.94 | 1.00 | 1.00 | 1.00 |
| | | | | 0.35 | 1.00 | 0.34 | 1.00 | 1.00 | 0.94 | 1.00 | 1.00 | 1.00 |
| | | | | 0.50 | 1.00 | 0.34 | 1.00 | 1.00 | 0.81 | 1.00 | 1.00 | 1.00 |
| | | | 12 | 0.15 | 1.00 | 1.00 | 1.00 | 1.00 | 0.98 | 1.00 | 1.00 | 1.00 |
| | | | | 0.35 | 1.00 | 0.34 | 1.00 | 1.00 | 0.81 | 1.00 | 1.00 | 1.00 |
| | | | | 0.50 | 1.00 | 0.34 | 1.00 | 1.00 | 0.62 | 1.00 | 1.00 | 1.00 |

Supplementary Table 4: Average clustering concordance $\overline{ARI}$ from compared methods for all simulation conditions (rows) with $K_{true} = 2$ underlying groups. Best performance within each row is colored in red.

| **n** | **LFC** | **pDE** | $\beta_0$ | $\phi_0$ | $\bar{K}^*_{FSC}$ | $\bar{K}^*_{iCl}$ | $\bar{K}^*_{HC}$ | $\bar{K}^*_{KM}$ | $\bar{K}^*_{NBMB}$ | $\bar{K}^*_{lMC}$ | $\bar{K}^*_{vMC}$ | $\bar{K}^*_{rMC}$ |
|---|---|---|---|---|---|---|---|---|---|---|---|---|
| 100 | 1.0 | 0.025 | 8 | 0.15 | 4.00 | 6.00 | 2.00 | 3.92 | 2.00 | 2.00 | 2.00 | 2.00 |
| | | | | 0.35 | 3.96 | 6.00 | 2.00 | 2.04 | 2.00 | 2.00 | 2.00 | 2.00 |
| | | | | 0.50 | 2.20 | 6.00 | 2.00 | 2.00 | 2.00 | 2.00 | 2.00 | 2.00 |
| | | | 12 | 0.15 | 4.08 | 6.00 | 2.00 | 3.84 | 2.00 | 2.00 | 2.00 | 2.00 |
| | | | | 0.35 | 3.96 | 6.00 | 2.00 | 2.00 | 2.00 | 2.00 | 2.00 | 2.00 |
| | | | | 0.50 | 2.32 | 6.00 | 2.00 | 2.00 | 2.00 | 2.00 | 2.00 | 2.00 |
| | | 0.050 | 8 | 0.15 | 4.08 | 6.00 | 2.00 | 4.00 | 2.00 | 2.00 | 2.00 | 2.00 |
| | | | | 0.35 | 3.96 | 6.00 | 2.00 | 2.80 | 2.00 | 2.00 | 2.00 | 2.00 |
| | | | | 0.50 | 3.92 | 6.00 | 2.00 | 2.36 | 2.00 | 2.00 | 2.00 | 2.00 |
| | | | 12 | 0.15 | 4.00 | 6.00 | 2.00 | 4.00 | 2.00 | 2.00 | 2.00 | 2.00 |
| | | | | 0.35 | 3.96 | 6.00 | 2.00 | 3.24 | 2.00 | 2.00 | 2.00 | 2.00 |
| | | | | 0.50 | 3.76 | 6.00 | 2.00 | 2.04 | 2.00 | 2.00 | 2.00 | 2.00 |
| | 2.0 | 0.025 | 8 | 0.15 | 4.12 | 6.00 | 2.00 | 4.00 | 2.00 | 2.00 | 2.00 | 2.00 |
| | | | | 0.35 | 4.00 | 6.00 | 2.00 | 4.00 | 2.00 | 2.00 | 2.00 | 2.00 |
| | | | | 0.50 | 4.00 | 6.00 | 2.00 | 3.88 | 2.00 | 2.00 | 2.00 | 2.00 |
| | | | 12 | 0.15 | 4.04 | 6.00 | 2.00 | 4.00 | 2.00 | 2.00 | 2.00 | 2.00 |
| | | | | 0.35 | 4.00 | 6.00 | 2.00 | 4.00 | 2.00 | 2.00 | 2.00 | 2.00 |
| | | | | 0.50 | 4.00 | 6.00 | 2.00 | 3.92 | 2.00 | 2.00 | 2.00 | 2.00 |
| | | 0.050 | 8 | 0.15 | 4.12 | 4.00 | 2.00 | 4.00 | 2.16 | 4.00 | 2.00 | 2.00 |
| | | | | 0.35 | 4.04 | 6.00 | 2.00 | 4.00 | 2.52 | 2.00 | 2.00 | 2.00 |
| | | | | 0.50 | 4.00 | 6.00 | 2.00 | 4.00 | 2.04 | 2.00 | 2.00 | 2.00 |
| | | | 12 | 0.15 | 4.00 | 4.00 | 2.00 | 4.00 | 2.44 | 4.00 | 2.00 | 2.00 |
| | | | | 0.35 | 4.08 | 6.00 | 2.00 | 4.00 | 2.48 | 2.00 | 2.00 | 2.00 |
| | | | | 0.50 | 4.04 | 6.00 | 2.00 | 4.00 | 2.00 | 2.00 | 2.00 | 2.00 |
| 200 | 1.0 | 0.025 | 8 | 0.15 | 4.28 | 6.00 | 2.00 | 4.00 | 2.48 | 2.00 | 2.00 | 2.00 |
| | | | | 0.35 | 4.12 | 6.00 | 2.00 | 2.00 | 2.00 | 2.00 | 2.00 | 2.00 |
| | | | | 0.50 | 4.08 | 6.00 | 2.00 | 2.00 | 2.00 | 2.00 | 2.00 | 2.00 |
| | | | 12 | 0.15 | 4.16 | 6.00 | 2.00 | 3.88 | 2.00 | 2.00 | 2.00 | 2.00 |
| | | | | 0.35 | 3.96 | 6.00 | 2.00 | 2.00 | 2.00 | 2.00 | 2.00 | 2.00 |
| | | | | 0.50 | 4.00 | 6.00 | 2.00 | 2.00 | 2.00 | 2.00 | 2.00 | 2.00 |
| | | 0.050 | 8 | 0.15 | 4.12 | 6.00 | 2.00 | 4.00 | 4.40 | 2.00 | 2.00 | 2.00 |
| | | | | 0.35 | 4.00 | 6.00 | 2.00 | 3.36 | 2.32 | 2.00 | 2.00 | 2.00 |
| | | | | 0.50 | 4.12 | 6.00 | 2.00 | 2.08 | 2.00 | 2.00 | 2.00 | 2.00 |
| | | | 12 | 0.15 | 4.28 | 6.00 | 2.00 | 4.00 | 3.40 | 2.00 | 2.00 | 2.00 |
| | | | | 0.35 | 4.00 | 6.00 | 2.00 | 3.56 | 2.16 | 2.00 | 2.00 | 2.00 |
| | | | | 0.50 | 4.00 | 6.00 | 2.00 | 2.12 | 2.00 | 2.00 | 2.00 | 2.00 |
| | 2.0 | 0.025 | 8 | 0.15 | 4.08 | 4.00 | 2.00 | 4.00 | 4.72 | 4.00 | 2.00 | 2.00 |
| | | | | 0.35 | 4.00 | 6.00 | 2.00 | 4.00 | 2.68 | 2.00 | 2.00 | 2.00 |
| | | | | 0.50 | 4.16 | 6.00 | 2.00 | 4.00 | 2.08 | 2.00 | 2.00 | 2.00 |
| | | | 12 | 0.15 | 4.08 | 4.00 | 2.00 | 4.00 | 4.44 | 4.00 | 2.00 | 2.00 |
| | | | | 0.35 | 4.16 | 6.00 | 2.00 | 4.00 | 2.68 | 2.00 | 2.00 | 2.00 |
| | | | | 0.50 | 4.16 | 6.00 | 2.00 | 3.92 | 2.04 | 2.00 | 2.00 | 2.00 |
| | | 0.050 | 8 | 0.15 | 4.04 | 4.00 | 2.00 | 4.00 | 2.52 | 4.00 | 4.00 | 4.00 |
| | | | | 0.35 | 4.28 | 5.12 | 2.00 | 4.00 | 2.16 | 3.92 | 2.00 | 2.00 |
| | | | | 0.50 | 4.00 | 6.00 | 2.00 | 4.00 | 2.16 | 2.00 | 2.00 | 2.00 |
| | | | 12 | 0.15 | 4.04 | 4.00 | 2.00 | 4.00 | 2.28 | 4.00 | 4.00 | 4.00 |
| | | | | 0.35 | 4.00 | 5.00 | 2.00 | 4.00 | 2.16 | 3.92 | 2.00 | 2.00 |
| | | | | 0.50 | 4.04 | 6.00 | 2.00 | 4.00 | 2.12 | 2.00 | 2.00 | 2.00 |

Supplementary Table 5: Average estimated orders $\bar{K}^*$ from compared methods for all simulation conditions (rows) with $K_{true} = 4$ underlying groups. Best performance within each row is colored in red.

| n | LFC | $p_{DE}$ | $\beta_0$ | $\phi_0$ | $\overline{ARI}_{FSC}$ | $\overline{ARI}_{iCl}$ | $\overline{ARI}_{HC}$ | $\overline{ARI}_{KM}$ | $\overline{ARI}_{NBMB}$ | $\overline{ARI}_{lMC}$ | $\overline{ARI}_{vMC}$ | $\overline{ARI}_{rMC}$ |
|---|---|---|---|---|---|---|---|---|---|---|---|---|
| 100 | 1.0 | 0.025 | 8 | 0.15 | 0.99 | 0.80 | 0.29 | 0.62 | 0.06 | 0.35 | 0.31 | 0.36 |
| | | | | 0.35 | 0.96 | 0.37 | 0.11 | 0.04 | 0.01 | 0.13 | 0.13 | 0.16 |
| | | | | 0.50 | 0.24 | 0.15 | 0.05 | 0.02 | 0.01 | 0.05 | 0.05 | 0.06 |
| | | | 12 | 0.15 | 0.98 | 0.78 | 0.34 | 0.65 | 0.07 | 0.36 | 0.33 | 0.36 |
| | | | | 0.35 | 0.93 | 0.44 | 0.11 | 0.04 | 0.01 | 0.15 | 0.13 | 0.15 |
| | | | | 0.50 | 0.24 | 0.14 | 0.03 | 0.02 | 0.01 | 0.06 | 0.05 | 0.06 |
| | | 0.050 | 8 | 0.15 | 0.99 | 0.80 | 0.37 | 0.98 | 0.26 | 0.38 | 0.38 | 0.38 |
| | | | | 0.35 | 0.99 | 0.76 | 0.37 | 0.26 | 0.06 | 0.35 | 0.34 | 0.33 |
| | | | | 0.50 | 0.98 | 0.59 | 0.26 | 0.09 | 0.03 | 0.21 | 0.24 | 0.24 |
| | | | 12 | 0.15 | 1.00 | 0.80 | 0.40 | 0.99 | 0.24 | 0.38 | 0.38 | 0.38 |
| | | | | 0.35 | 0.99 | 0.77 | 0.34 | 0.33 | 0.05 | 0.35 | 0.35 | 0.36 |
| | | | | 0.50 | 0.94 | 0.62 | 0.28 | 0.06 | 0.03 | 0.23 | 0.23 | 0.25 |
| | 2.0 | 0.025 | 8 | 0.15 | 0.99 | 0.78 | 0.38 | 1.00 | 0.26 | 0.36 | 0.37 | 0.37 |
| | | | | 0.35 | 1.00 | 0.78 | 0.37 | 0.93 | 0.11 | 0.38 | 0.38 | 0.37 |
| | | | | 0.50 | 1.00 | 0.80 | 0.33 | 0.62 | 0.11 | 0.37 | 0.34 | 0.38 |
| | | | 12 | 0.15 | 1.00 | 0.78 | 0.40 | 1.00 | 0.21 | 0.37 | 0.39 | 0.37 |
| | | | | 0.35 | 1.00 | 0.79 | 0.40 | 0.94 | 0.10 | 0.38 | 0.36 | 0.37 |
| | | | | 0.50 | 1.00 | 0.80 | 0.33 | 0.69 | 0.10 | 0.35 | 0.34 | 0.37 |
| | | 0.050 | 8 | 0.15 | 1.00 | 1.00 | 0.39 | 1.00 | 0.41 | 1.00 | 0.38 | 0.37 |
| | | | | 0.35 | 1.00 | 0.79 | 0.38 | 1.00 | 0.33 | 0.39 | 0.39 | 0.39 |
| | | | | 0.50 | 1.00 | 0.79 | 0.39 | 0.99 | 0.33 | 0.38 | 0.38 | 0.38 |
| | | | 12 | 0.15 | 1.00 | 1.00 | 0.39 | 1.00 | 0.37 | 1.00 | 0.39 | 0.38 |
| | | | | 0.35 | 0.99 | 0.78 | 0.42 | 1.00 | 0.34 | 0.39 | 0.40 | 0.40 |
| | | | | 0.50 | 1.00 | 0.79 | 0.40 | 0.99 | 0.35 | 0.39 | 0.39 | 0.39 |
| 200 | 1.0 | 0.025 | 8 | 0.15 | 0.98 | 0.78 | 0.34 | 0.68 | 0.24 | 0.38 | 0.36 | 0.36 |
| | | | | 0.35 | 0.98 | 0.76 | 0.16 | 0.05 | 0.06 | 0.16 | 0.16 | 0.19 |
| | | | | 0.50 | 0.99 | 0.45 | 0.06 | 0.02 | 0.03 | 0.07 | 0.07 | 0.09 |
| | | | 12 | 0.15 | 0.99 | 0.80 | 0.32 | 0.67 | 0.30 | 0.36 | 0.34 | 0.37 |
| | | | | 0.35 | 0.99 | 0.76 | 0.16 | 0.05 | 0.08 | 0.18 | 0.15 | 0.20 |
| | | | | 0.50 | 0.98 | 0.42 | 0.05 | 0.02 | 0.03 | 0.05 | 0.07 | 0.08 |
| | | 0.050 | 8 | 0.15 | 1.00 | 0.80 | 0.41 | 0.99 | 0.41 | 0.36 | 0.38 | 0.36 |
| | | | | 0.35 | 1.00 | 0.79 | 0.34 | 0.35 | 0.28 | 0.35 | 0.35 | 0.35 |
| | | | | 0.50 | 1.00 | 0.79 | 0.30 | 0.08 | 0.14 | 0.22 | 0.24 | 0.24 |
| | | | 12 | 0.15 | 0.98 | 0.80 | 0.40 | 0.99 | 0.39 | 0.38 | 0.37 | 0.38 |
| | | | | 0.35 | 1.00 | 0.80 | 0.36 | 0.38 | 0.29 | 0.36 | 0.35 | 0.37 |
| | | | | 0.50 | 1.00 | 0.79 | 0.28 | 0.08 | 0.13 | 0.24 | 0.24 | 0.28 |
| | 2.0 | 0.025 | 8 | 0.15 | 0.99 | 1.00 | 0.39 | 1.00 | 0.34 | 1.00 | 0.36 | 0.35 |
| | | | | 0.35 | 1.00 | 0.81 | 0.37 | 0.96 | 0.36 | 0.36 | 0.37 | 0.34 |
| | | | | 0.50 | 0.99 | 0.80 | 0.34 | 0.72 | 0.34 | 0.35 | 0.37 | 0.36 |
| | | | 12 | 0.15 | 0.99 | 1.00 | 0.39 | 1.00 | 0.32 | 1.00 | 0.36 | 0.35 |
| | | | | 0.35 | 1.00 | 0.80 | 0.40 | 0.95 | 0.40 | 0.36 | 0.35 | 0.35 |
| | | | | 0.50 | 0.99 | 0.81 | 0.36 | 0.68 | 0.32 | 0.37 | 0.36 | 0.36 |
| | | 0.050 | 8 | 0.15 | 1.00 | 1.00 | 0.43 | 1.00 | 0.45 | 1.00 | 1.00 | 1.00 |
| | | | | 0.35 | 0.99 | 0.89 | 0.42 | 1.00 | 0.44 | 0.98 | 0.36 | 0.36 |
| | | | | 0.50 | 1.00 | 0.80 | 0.39 | 0.99 | 0.47 | 0.37 | 0.37 | 0.37 |
| | | | 12 | 0.15 | 1.00 | 1.00 | 0.38 | 1.00 | 0.51 | 1.00 | 1.00 | 1.00 |
| | | | | 0.35 | 1.00 | 0.90 | 0.39 | 1.00 | 0.44 | 0.98 | 0.37 | 0.37 |
| | | | | 0.50 | 1.00 | 0.79 | 0.42 | 0.99 | 0.43 | 0.37 | 0.37 | 0.37 |

Supplementary Table 6: Average clustering concordance $\overline{ARI}$ from compared methods for all simulation conditions (rows) with $K_{true} = 4$ underlying groups. Best performance within each row is colored in red.

### Section D: TCGA Breast Cancer Analyses

#### D1: Pre-processing

The RNA-Seq gene expression quantification dataset and annotations were acquired from the Illumina Hiseq platform using the *GDCquery()* function from the `TCGAbiolinks` package (Colaprico et al., 2015). FPKM values for this dataset were also acquired using the *GDCquery()* function. We first normalized the data for differences in sequencing depth using `DESeq2` (Love et al., 2014). Then, we pre-filtered out genes with median FPKM $\leq 1$. From there, we pre-filtered out samples whose ABSOLUTE purity estimates were $< 0.9$. ABSOLUTE purity estimates were obtained from Aran et al. (2015). From this subset, we pre-filtered out genes with median normalized count $< 500$ or with median absolute deviation (MAD) below the $50^{th}$ quantile. The plate information we used as batch effects was obtained from the MD Anderson TCGA Batch Effects Viewer, using the 'current' version of the GDC Index file of TCGA BRCA RNASeq-Counts. Plates with only one sample were merged together as one plate for proper analysis.

#### D2: Heatmap of cluster-discriminatory genes

In this section, we show a heatmap of all cluster-discriminatory genes derived by $FSC_{adj}$. Our results showed good performance across all methods except `iCluster+` in discriminating between Basal and Luminal subtypes, likely because `iCluster+` does not take into account the overdispersed nature of the counts. HC and KM did not detect differences between Luminal A and Luminal B subtypes, clustering them into one group. Of all the methods, $FSC_{adj}$ achieved the best concordance with the annotated subtypes.

#### D3: Gene set enrichment analysis

We performed gene set enrichment analysis on the cluster-discriminatory genes discovered via $FSC$ and $FSC_{adj}$. Figure 4 shows the top 10 enriched biological processes and pathways found using the `TCGAbiolinks` R package (Colaprico et al., 2015) on the list of cluster-discriminatory genes from $FSC$ and $FSC_{adj}$. In either case, we find that proportions of genes are high for enriched pathways related to the cell cycle. Enriched pathways included those related to the cell cycle (e.g. 'Cell Cycle Control of Chromosomal Replication') and signaling pathways (e.g. 'Aryl Hydrocarbon Receptor Signaling'), and other 'Molecular Mechanisms of Cancer'. Figure 5 shows the top 10 overlapping MSigDB gene sets (Mootha et al., 2003; Subramanian et al., 2005) with the list of cluster-discriminatory genes from $FSC_{adj}$ that were up-regulated and down-regulated in basal samples ($basalUP$ and $basalDOWN$, respectively). Top overlapping gene sets pertain to genes that have either been associated with or been known to delineate subtypes of breast cancer. The top overlapping gene set for $basalUP$ and $basalDOWN$ are 'SMID_BREAST_CANCER_BASAL_UP' and 'SMID_BREAST_CANCER_BASAL_DN' (Smid et al., 2008), respectively, validating the subtype-delineating nature of the cluster-discriminatory genes found by $FSC_{adj}$. Some other top overlapping gene sets included the Vantveer and Charafe breast cancer gene set

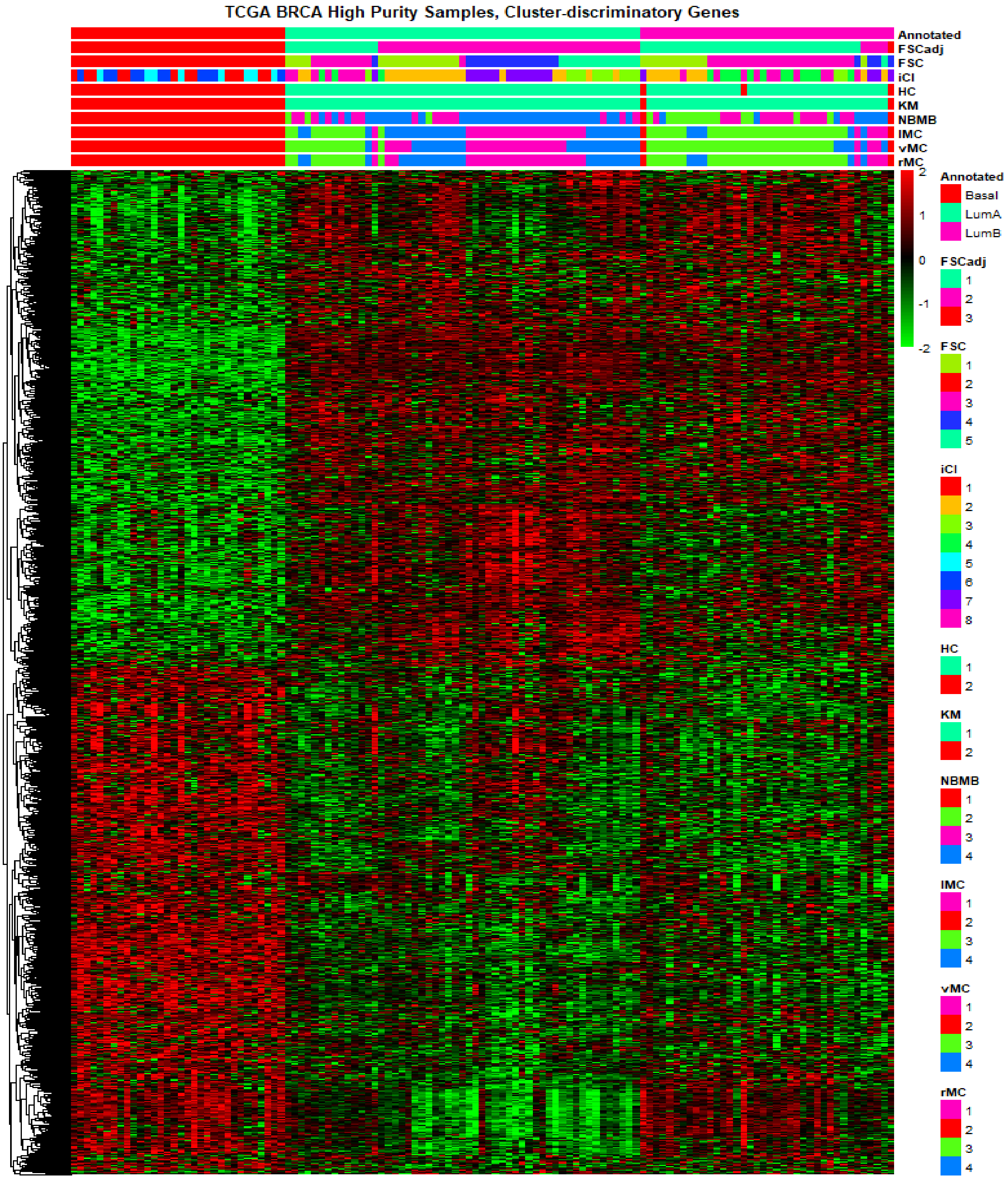


Supplementary Figure 3: Heatmap of the cluster-discriminatory genes discovered via $FSC_{adj}$ analysis, with column annotations for clustering labels (top). Column ordering is based on annotated subtypes, and samples are ordered within subtypes by decreasing order of maximum posterior probability via $FSC_{adj}$. All clustering labels except `iCluster+` distinguish well between Basal and Luminal subtypes, but $FSC_{adj}$ best distinguishes between Luminal A and Luminal B samples.

modules (van 't Veer et al., 2002; Charafe-Jauffret et al., 2005). Similar gene sets were discovered when all cluster-discriminatory genes were analyzed jointly.

#### D4: Analysis without purity filtering

We processed the dataset the same way as in Section D1, but without filtering samples based upon ABSOLUTE purity estimates. We narrowed down to the samples found in the (Koboldt et al., 2012) study for head-to-head comparison. We performed all methods on the resulting dataset of 581 samples and 4261 genes. This is slightly different from what Koboldt et al. (2012) cited (547 samples), but this is because their supplementary file annotated each sample by the patient ID rather than the unique barcode. Thus, we found an additional 34 samples, which were likely gathered from the same patients after the study was published. We assumed here that multiple samples from the same patient corresponded to the same tumor type.

| | $K^*$ (5) | ARI (1) | NMI (1) | NVI (0) | NID (0) |
|---|---|---|---|---|---|
| anno | 5 | | | | |
| $FSC$ | 7 | 0.316 | 0.369 | 0.732 | 0.631 |
| $FSC_{adj}$ | 8 | 0.245 | 0.295 | 0.78 | 0.705 |
| iCl | 6 | 0.257 | 0.28 | 0.805 | 0.72 |
| HC | 2 | 0.004 | 0.003 | 0.997 | 0.997 |
| KM | 2 | 0.348 | 0.293 | 0.731 | 0.707 |
| NBMB | 15 | -0.008 | 0.097 | 0.932 | 0.903 |
| lMC | 15 | 0.157 | 0.259 | 0.788 | 0.741 |
| vMC | 12 | 0.164 | 0.268 | 0.786 | 0.732 |
| rMC | 12 | 0.169 | 0.274 | 0.781 | 0.726 |
| SigI | 14 | 0.258 | 0.333 | 0.73 | 0.667 |
| SigU | 13 | 0.272 | 0.35 | 0.736 | 0.65 |

Supplementary Table 7: Selected order ($K^*$) and clustering concordance between compared methods and annotated TCGA Breast Cancer subtypes. FSCseq was run with adjustment ($FSC_{adj}$) and without adjustment ($FSC$) for plate effect, and each of the clustering labels were compared to annotated subtypes (anno). Results from unsupervised (SigU) and semi-supervised (SigI) clustering from Koboldt et al. (2012) are also shown. For each column, the best performing metric is colored in red. The value in parentheses in the column headings represent optimal values. For ARI and NMI, values close to 1 indicate good clustering, and values close to 0 indicate poor clustering. For NVI and NID, values close to 0 indicate good clustering, and values close to 1 indicate poor clustering.

Supplementary Table 7 shows the clustering performance of compared methods, and Supplementary Figure 6 shows a heatmap of the genes that were found to be discriminatory via $FSC_{adj}$. SigU and SigI represent unsupervised and semi-supervised clustering labels derived from Koboldt et al. (2012). SigI was derived from a narrowed-down intrinsic list of significant genes. Most methods performed well in distinguishing the basal subtype, except for SigU. Of the methods, $FSC$ attained best performance in 2 of the 4 clustering metrics: $NMI$ and $NID$. KM attained the highest $ARI$, but KM only delineated between basal vs. rest, and failed to distinguish samples across the other 4 subtypes. iCluster+ selected the order closest to the truth, but $FSC$ yielded better cluster concordance.

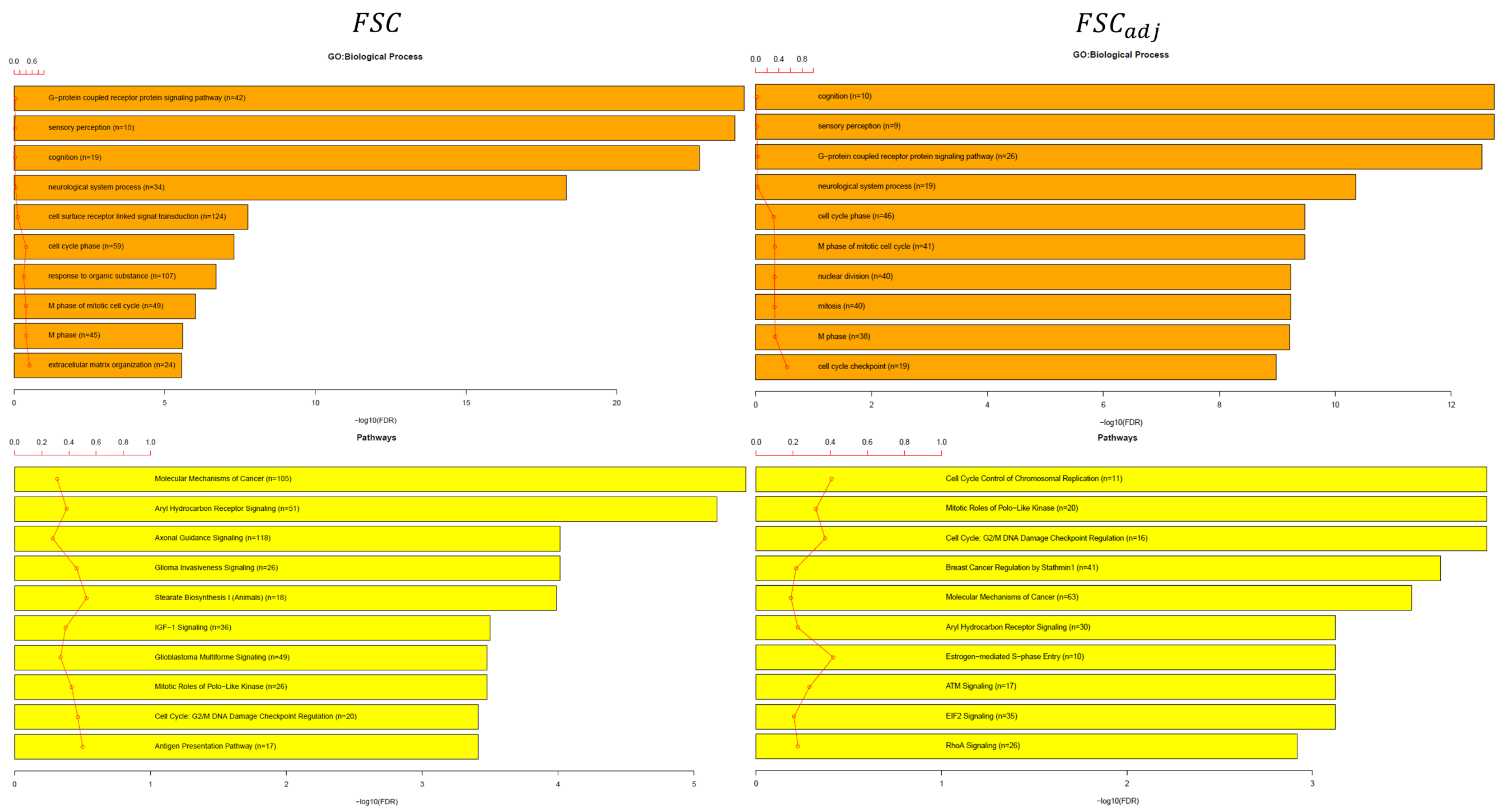


Supplementary Figure 4: Gene ontology biological processes and enriched pathways of cluster-discriminatory genes found by $FSC$ (left) and $FSC_{adj}$ (right). The bars represent p-value corrected FDR (in $-\log$ scale), and the red lines represent ratios of list genes found in each pathway over the total number of genes in that pathway.

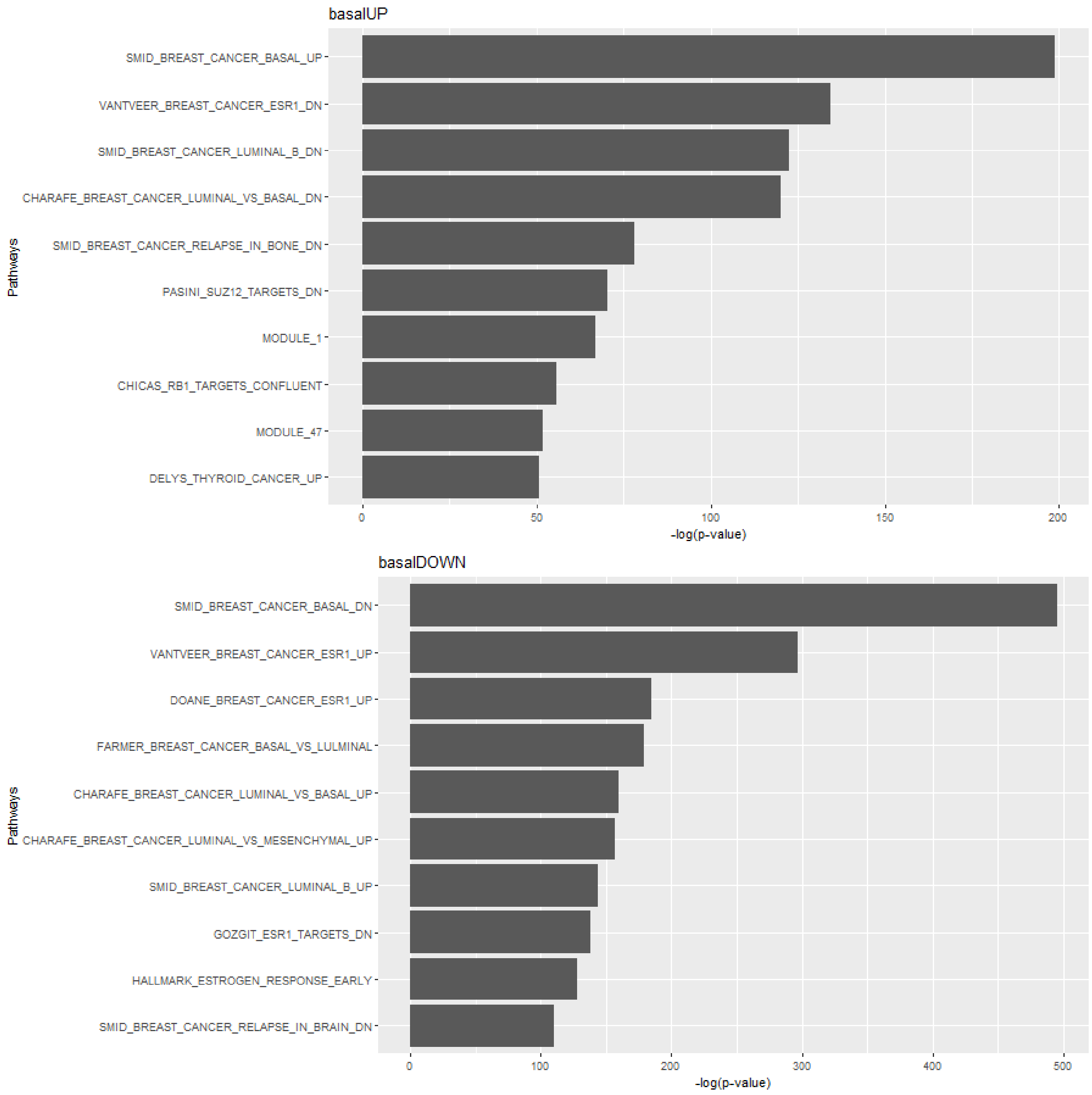


Supplementary Figure 5: GSEA analyses on $basalUP$ (basal up-regulated, top) and $basalDOWN$ (basal down-regulated, bottom) subsets of cluster-discriminatory genes discovered from $FSC_{adj}$. Only the top five overlapping gene sets shown.

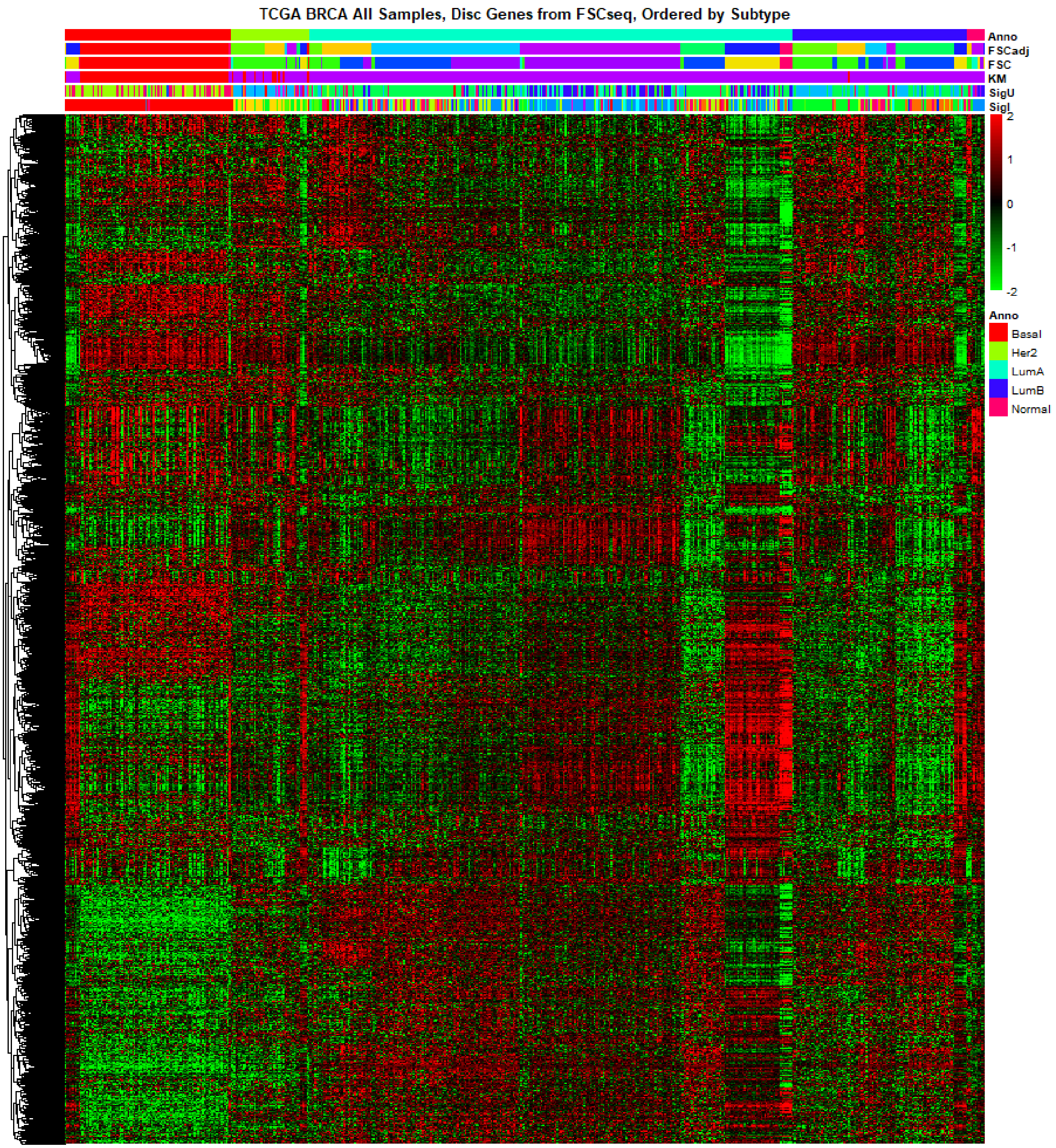


Supplementary Figure 6: Heatmap of the genes found to be discriminatory via $FSC_{adj}$, ordered by annotated subtypes (Anno). Compared clustering labels were from FSCseq with ($FSC_{adj}$) and without ($FSC$) adjustment for plate, K-medoids (KM), and unsupervised (SigU) and semi-supervised clustering based on intrinsic gene list (SigI) (Koboldt et al., 2012).

Interestingly, cluster labels from $FSC_{adj}$ were less concordant with annotated subtypes than those from $FSC$. This is because of differences in the purity of tumor samples, causing heterogeneity that likely confounded the effects from plate. The heatmap of cluster-discriminatory in Supplementary Figure 6 shows this heterogeneity within samples of the annotated subtypes, which $FSC_{adj}$ and $FSC$ delineate, although the annotated groups clustered such heterogeneous samples together. Still, $FSC_{adj}$ and $FSC$ selected $K^*$ that was much closer to the number of subtypes than previous results by SigU and SigI, which suggest that our clusters were more closely related to the subtypes, although similarly confounded by the sample heterogeneity.
